# Hypertrophic cardiomyopathy ß-cardiac myosin mutation (P710R) leads to hypercontractility by disrupting super-relaxed state

**DOI:** 10.1101/2020.11.10.375493

**Authors:** Alison Schroer Vander Roest, Chao Liu, Makenna M Morck, Kristina Bezold Kooiker, Gwanghyun Jung, Dan Song, Aminah Dawood, Arnav Jhingran, Gaspard Pardon, Sara Ranjbarvaziri, Giovanni Fajardo, Mingming Zhao, Kenneth S Campbell, Beth L Pruitt, James A Spudich, Kathleen M Ruppel, Daniel Bernstein

## Abstract

Hypertrophic cardiomyopathy (HCM) is the most common inherited form of heart disease, associated with over 1000 mutations, many in β-cardiac myosin (MYH7). Molecular studies of myosin with different HCM mutations have revealed a diversity of effects on ATPase and load-sensitive rate of detachment from actin. It has been difficult to predict how such diverse molecular effects combine to influence forces at the cellular level and further influence cellular phenotypes. This study focused on the P710R mutation that dramatically decreased in vitro motility velocity and actin-activated ATPase, in contrast to other MYH7 mutations. Optical trap measurements of single myosin molecules revealed that this mutation reduced the step size of the myosin motor and the load sensitivity of the actin detachment rate. Conversely, this mutation destabilized the super-relaxed state in longer, two-headed myosin constructs, freeing more heads to generate force. Micropatterned hiPSC-cardiomyocytes CRISPR-edited with the P710R mutation produced significantly increased force (measured by traction force microscopy) compared with isogenic control cells. The P710R mutation also caused cardiomyocyte hypertrophy and cytoskeletal remodeling as measured by immunostaining and electron microscopy. Cellular hypertrophy was prevented in the P710R cells by inhibition of ERK or Akt. Finally, we used a computational model that integrated the measured molecular changes to predict the measured traction forces. These results confirm a key role for regulation of the super-relaxed state in driving hypercontractility in HCM with the P710R mutation and demonstrate the value of a multiscale approach in revealing key mechanisms of disease.

**Significance Statement:** Heart disease is the leading cause of death worldwide, and hypertrophic cardiomyopathy (HCM) is the most common inherited form of heart disease, affecting over 1 in 200 people. Mutations in myosin, the motor protein responsible for contraction of the heart, are a common cause of HCM but have diverse effects on the biomechanics of the myosin protein. We demonstrate that complex biomechanical effects of mutations associated with heart disease can be effectively studied and understood using a multi-scale experimental and computational modeling approach. This work confirmed an important role for disruption of the super-relaxed state for one particular HCM mutation, and our approach can be extended to aid in the development of new targeted therapies for patients with different mutations.

## Introduction

Hypertrophic cardiomyopathy (HCM) is one of the most prevalent genetic diseases of the heart, affecting over 1 in 200 individuals (1, 2), and is a leading cause of sudden cardiac death (3). HCM is characterized by cardiomyocyte hypertrophy, myofibril disarray, hypercontractility, and diastolic dysfunction. Tissue remodeling, including interstitial fibrosis, can eventually progress to heart failure and death (4–6). Over 1000 causative mutations have been identified, with the majority in genes encoding sarcomeric proteins responsible for generating and regulating contraction. Roughly a third of mutations are located in β-cardiac myosin, the primary ventricular motor protein in humans (7).

With the centrality of hypercontractility in disease phenotypes, HCM mutations were hypothesized to increase the activity of myosin at the protein level, resulting in increased force production at the sarcomere and cellular levels that propagates to the whole organ level (8). Myosin protein activity is characterized by biochemical and biophysical measurements. The activity of actively-cycling myosin interacting with actin is characterized by the rate of ATP turnover, the rate of detachment from actin, force production, step size, and actin-sliding velocity. Myosin that is not actively cycling resides in a super-relaxed state (SRX) (9), associated with a folded-back conformation not available for actin interaction. Biochemical and biophysical assessments of HCM mutations in purified human β-cardiac myosin have revealed various changes in both actively-cycling myosin (10–14) and SRX proportions (15–18).

Myosin’s converter domain, or hinge responsible for the powerstroke, is a hotspot for pathogenic mutations (19–22). The P710 residue is located at the proximal edge of the converter domain, and at least three HCM mutations have been identified at this site, including P710R (23–25). A previous report on the P710R mutation found that actively cycling heads have lower ATP turnover activity and duty ratio compared to both WT myosin and other HCM mutations (26). The discrepancy between this reduced function in actively cycling myosin and the hypercontractile clinical phenotype invites a comprehensive, multiscale assessment of the effects of the P710R mutation on myosin activity and SRX proportions.

In the past, it has been difficult to determine early mechanisms of HCM disease triggered by mutations in β-MHC because of the inability to culture human cardiomyocytes. Samples from hearts obtained at the time of transplant reflect a combination of primary and secondary pathologies. Rodents fail to recapitulate many human heart diseases, and their adult ventricles predominantly express the α-MHC isoform, which has different kinetics from β-MHC (27–29). However, the expansion of CRISPR/Cas-9 protocols for human induced pluripotent derived stem cell (hiPSC) gene editing coupled with efficient differentiation into cardiomyocytes (hiPSC-CMs) provides a valuable model system for studying early mechanisms of disease in a controlled context (28). In the past, hiPSC-CM models have been limited by the relative immaturity of the cardiomyocytes and high population heterogeneity (30). In traditional two-dimensional culture, these cells have disorganized myofibrils, impaired calcium handling, and immature cell-signaling (31, 32). However, recent advances in microengineered environments can provide external environmental cues that better recapitulate *in vivo* conditions to promote myofibril alignment and accelerate maturation of both contractile machinery and cell signaling (32, 33). We have previously developed a hydrogel platform with rectangles of extracellular matrix (ECM) at a 7:1 aspect ratio (similar to that of cardiomyocytes in the left ventricle) that, combined with traction force microscopy (TFM), allows for single-cell assessment of both cellular organization and biomechanics (32–34). When combined with measurements of myosin function at the molecular level, these cellular measurements can provide validation of the molecular basis of force generation and resultant disease mechanisms of HCM.

Finally, computational models can provide key insights into the interactions between related dynamic parameters and the resultant implications for the total production of force. Computational models incorporating different degrees of detail and different components of sarcomere structure have been used for decades to answer questions relating to fundamental muscle mechanics (35–44) and alterations to thin filament regulation in the context of cardiac disease and HCM (45, 46). A new model of myosin activity which incorporates an OFF state representative of the SRX state has recently been validated against experimental measurements of cardiac muscle (47). This study concluded that force-sensitive regulation of the OFF state significantly improves the fit to experimental data, but this model has not previously been used in the context of MHY7 mutations nor hiPSC-CMs (47).

In this work, we used multi-scale experimental techniques to assess the biomechanical effects of the HCM mutation P710R which demonstrates decreased activity at the level of the motor domain yet increased force generation at the cell level. We furthermore used a computational model to integrate the molecular findings and found that altered regulation of the SRX state is an essential driver of hypercontractility for this mutation. Our multi-scale experimental findings combined with computational modeling enabled us to assess the relative contributions of individual molecular parameters to cellular contraction.

## Results

### The P710R mutation reduced load sensitivity and step size of single myosin molecules

We first focused on the motor domain of myosin by using the recombinant human β-cardiac myosin sS1 domain (catalytic domain plus the essential light chain binding portion of the lever arm). The properties of individual myosin heads interacting with actin were assessed by optical trapping using harmonic force spectroscopy (HFS). In this technique, the durations of binding events between a single myosin and an actin filament under different load forces are measured in physiological (2 mM) ATP conditions. The sample stage oscillates sinusoidally so that by the randomness of where myosin initially attaches to actin, a range of mean forces are automatically applied over the course of many binding events (48). This technique has been used to quantify changes in the load-dependent detachment rate of β-cardiac myosin due to mutations and myosin inhibitor and activators (12).

The detachment rate is an exponential function of the mean load force (*F*):

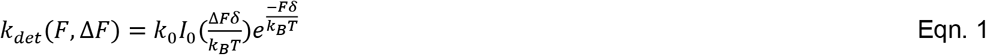

where *k*_0_ is the rate at zero load, *δ* is the distance to the transition state of the rate-limiting step in the bound state (a measure of force sensitivity), *k*_B_ is the Boltzmann constant, *T* is temperature, and *I*_0_ is the zeroth-order modified Bessel function of the first kind (to correct for the harmonic force with amplitude *ΔF*). The release of ADP is the rate limiting step for detachment from actin at high (2 mM) ATP concentrations, and this step is sensitive to load force. Thus, the detachment rate *k*_0_ and its load sensitivity *δ* determined by HFS correspond to the rate of ADP release and its load sensitivity, respectively. WT β-cardiac myosin had detachment rate at zero load *k*_0_ = 104 ± 10 s^−1^ and force sensitivity *δ* = 1.39 ± 0.06 nm, consistent with previous results (12, 48–50). The unloaded detachment rate of P710R was not significantly changed (87 ± 5 s^−1^, p = 0.12), while *δ* was dramatically reduced (0.31 ± 0.03 nm, p < 0.0001) (**Table 1** and **Figure 1AB**).

**Table 1:** Parameters measured for β-cardiac myosin sS1. The rate of detachment from actin at zero load *k*_0_, its force sensitivity *δ* and myosin’s step size *d* were measured from single molecules using the HFS technique. *k*_cat_ and *K*_app_ were measured using a colorimetric actin-activated ATPase assay and were previously reported (26). Unloaded motility velocities *V* were measured by the motility assay with actin filaments. Calcium sensitivity parameters *pCa*_50_ and Hill coefficient *n* were measured by motility assay with regulated thin filaments. Data are presented as mean ± standard error of the mean. * indicates p < 0.05 and **** indicates p < 0.0001 different between P710R and WT.

| | $k_0$ (s<sup>-1</sup>) | $\delta$ (nm) | $d$ (nm) | $k_{cat}$ (s<sup>-1</sup>) | $K_{app}$ (μM) | $V$ (nm s<sup>-1</sup>) | $pCa_{50}$ | $n$ |
| --- | --- | --- | --- | --- | --- | --- | --- | --- |
| **WT** | 104 ± 10 | 1.39 ± 0.06 | 5.2 ± 0.3 | 4.1 ± 0.4 | 33 ± 6 | 762 ± 16 | 6.46 ± 0.04 | 2.2 ± 0.2 |
| **P710R** | 87 ± 5 | 0.31 ± 0.03**** | 1.9 ± 0.3**** | 2.50 ± 0.1* | 14 ± 1* | 239 ± 9**** | 6.48 ± 0.07 | 3.1 ± 0.6 |

**Figure 1:**
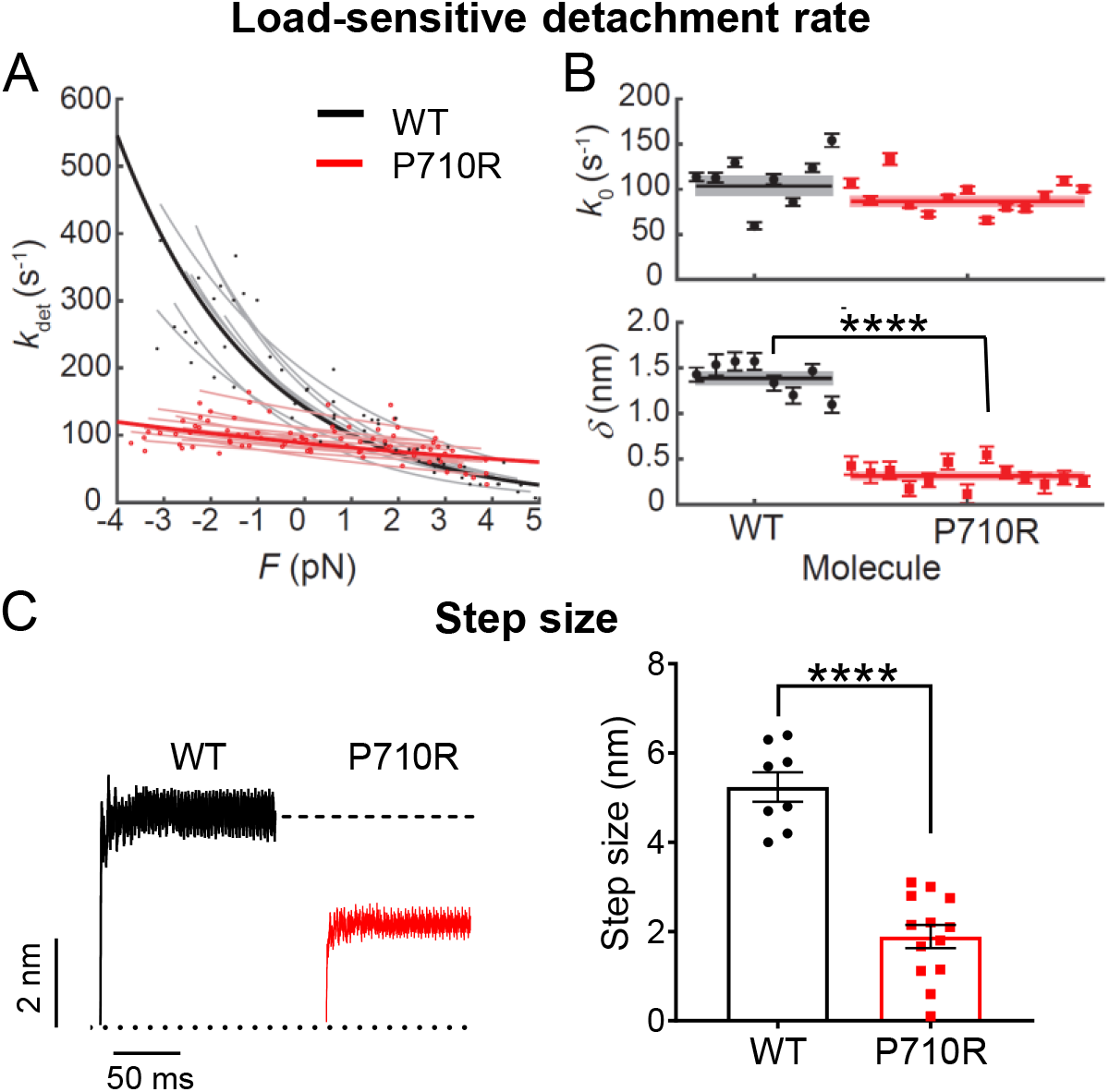
Single molecules of β-cardiac myosin sS1 with P710R mutation had reduced load sensitivity and step size. **(A)** Measurements of myosin’s load sensitive rate of detachment from actin *k*_det_(*F*) using the harmonic force spectroscopy technique in a dual-beam optical trap. Positive forces represent load in the opposite direction of the powerstroke (resistive), and negative forces represent load in the same direction of the powerstroke (assistive). Each light line is a fit of Eqn. 1 to data from one molecule, each with a few hundred binding events. **(B)** The fitted parameters *k*_0_ (rate at zero load) and *δ* (load sensitivity) of each molecule corresponding to light lines in (A). Error bars represent the error in the parameter fit for each molecule. Horizontal lines represent weighted means across all molecules, and shaded rectangles represent s.e.m. (**C**) Averaged start-aligned position traces of binding events from two example molecules revealing the power stroke of myosin, which occur within milliseconds of actin binding. Step size values of multiple molecules are shown on the right. Error bars represent s.e.m. Values are given in Table 1. See also methods and Figure S1. **** indicates p < 0.0001.

Further analysis of the average trapped bead displacement during binding events revealed that the mutation decreased myosin’s step size by ~60% (WT *d* = 5.2 ± 0.3 nm, P710R *d* = 1.9 ± 0.3 nm, p < 0.0001) (**Table 1, Figure 1CD, Figure S1**).

### The P710R mutation reduced *k*_cat_, duty ratio, and velocity but not calcium sensitivity of myosin in ensemble

We next assessed the effects of the P710R mutation on properties of myosin sS1 in ensemble. Myosins with the P710R mutation had significantly reduced actin-activated ATPase activity (WT *k*_cat_ = 4.1 ± 0.4 s^−1^, P710R *k*_cat_ = 2.5 ± 0.1 s^−1^) and higher apparent actin affinity (WT *K*_app_ = 33 ± 6 μM, P710R *K*_app_ = 14 ± 1 μM) as reported previously (26) (**Table 1)**. Given the single molecule and ensemble ATPase measurements, the duty ratio can be estimated as a function of load force (*F*) (12):

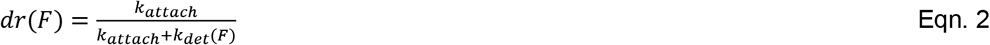

where the attachment rate *k*_attach_, assumed to be independent of force, is given by

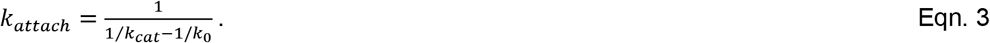

Our calculations suggest that P710R has a much lower duty ratio than WT myosin at low or resistive (positive) loads (**Figure 2A**), consistent with a previously reported prediction (26). Since duty ratio is the fraction of heads bound to actin in a force-producing state at any given time, P710R is predicted to generate less force per head on average than WT at those loads. Our calculations predict that P710R has a slightly higher duty ratio than WT under high assistive (negative) loads, suggesting that P710R heads may not release actin as efficiently in an actively shortening sarcomere.

**Figure 2:**
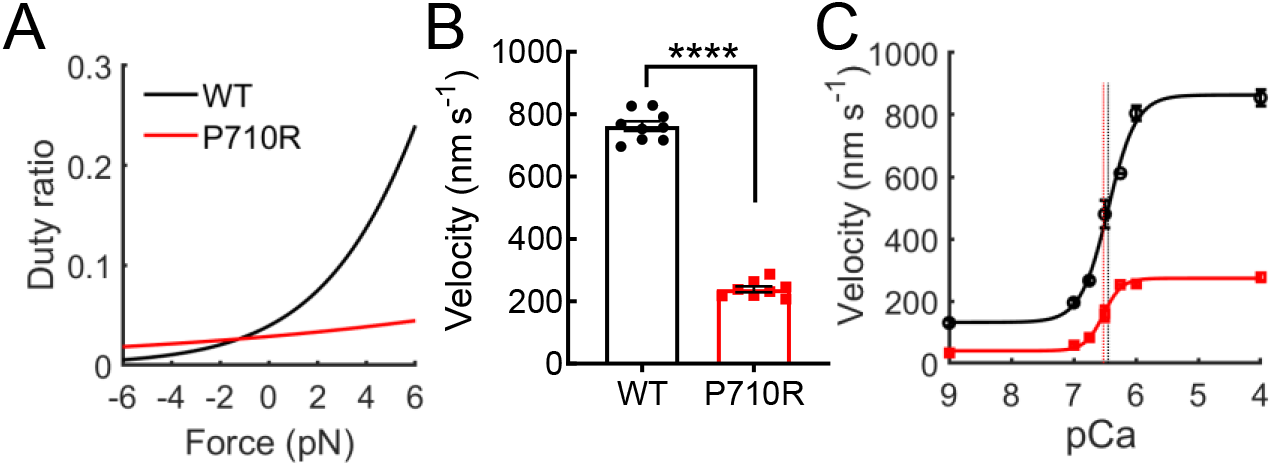
β-cardiac myosin sS1 ensembles with P710R mutation had altered duty ratio, reduced actin sliding velocity, and unchanged calcium sensitivity. (**A**) Calculated duty ratio across load forces based on ATPase and actin detachment rate measurements. (**B**) Actin sliding velocity (mean velocity including stuck filaments, MVIS; see methods) in the unloaded motility assay. Each data point represents one independent experiment. (**C**) Velocities (MVIS) of regulated thin filaments at various calcium concentrations measured by the motility assay. pCa = −log_10_[Ca]. Curves are fits to the Hill equation of averaged data from multiple independent experiments, and vertical dashed lines represent the fitted *pCa*_50_’s. Error bars represent s.e.m. Values are given in Table 1. See also Figure S2. **** indicates p < 0.0001.

Actin sliding velocity in an unloaded motility assay was reduced by ~60% for P710R compared to WT (WT *V* = 762 ± 16 nm/s, P710R *V* = 239 ± 9 nm/s, p < 0.0001) (**Table 1, Figure 2B**, Figure S2, Movies S1 and S2). Velocity can be approximated as the step size divided by the actin-bound time, which is inversely proportional to the actin detachment rate: *V = d/t*_bound_ = *d*k*_det_. Thus, the 60% reduction in unloaded motility velocity can be explained by the 60% reduction in P710R’s step size and an unchanged unloaded detachment rate *k*_0_. A previous study of this mutation failed to predict the measured decrease in velocity likely because the authors did not account for a major reduction in step size (26). We further observed that 5-15% of actin filaments were immobile for P710R compared to 0-5% for WT (Figure S2). This suggests that the mutation may induce a less proper or stable protein fold in a small percentage of myosin molecules which, as a result, have reduced function in moving actin (see also methods and discussion).

In addition to determining motility velocity, myosin’s actin-detachment kinetics may affect the calcium sensitivity of sarcomeres through the activation of the thin filament by bound myosin heads (44). In the thin filament, binding sites on actin are blocked by the regulatory proteins troponin and tropomyosin when calcium levels are low (in diastole) and are activated and accessible to myosin heads when calcium levels rise (in systole). However, heads that stay bound to actin longer may cooperatively activate the thin filament at lower calcium concentrations (51, 52), thereby altering the calcium sensitivity. While P710R had an unchanged unloaded actin-detachment rate *k*_0_, the effect of its reduced force sensitivity *δ* on this mechanism of calcium sensitivity is unclear. To this end, we measured the sliding velocities of regulated thin filaments (RTF, actin decorated with troponin and tropomyosin) at various calcium concentrations in the motility assay. RTF velocities increased as calcium concentration increased through the physiological range (pCa7-6, or 100 nM to 1 μM), and velocities saturated at high calcium levels (pCa4) (**Figure 2C**). We found that the P710R mutation did not significantly affect the calcium sensitivity; neither the calcium concentration for half-maximum velocity (P710R *pCa*_50_ = 6.48 ± 0.07, WT *pCa*_50_ = 6.46 ± 0.04, p = 0.78) nor the Hill coefficient (P710R *n* = 3.1 ± 0.6, WT *n* = 2.2 ± 0.2, p = 0.23) were significantly different from WT (**Table 1**; **Figure 2C**).

Taken together, the above measurements of the myosin motor domain at the single molecule and ensemble levels do not suggest a clear mechanism of hypercontractility by the HCM mutation P710R. In fact, they appear to suggest hypocontractility, in conflict with the mutation’s physiological effects in patients.

### The P710R mutation disrupted the SRX state of myosin in vitro

We and others have previously shown that the ability of myosin to form the folded-back interacting-heads motif structure is critical for regulating myosin activity, and many HCM-causing mutations appear to disrupt the ability of myosin to enter this state (15–18, 53). To assess the ability of myosin to form this folded-back “off” state in vitro, we used 2-headed myosin constructs as previously described (18). We measured a >40% reduction in the actin-activated ATPase rate (*k*_cat_) of a long-tailed (25-hep) WT myosin construct compared to a short-tailed (2-hep) WT construct (2-hep *k*_cat_ = 7.2 ± 0.7 s^−1^, 25-hep *k*_cat_ = 4.1 ± 0.3 s^−1^) (**Figure 3A,B**), consistent with our previous observations (15, 17, 18). This difference can be attributed to the sequestration of a large fraction of the myosin in the folded-back state in the long-tailed population, thus preventing the heads from binding to actin and entering the ATPase cycle. In contrast, we found no significant difference between the ATPase rate of the P710R short- and long-tailed myosins (P710R 2 hep *k*_cat_ = 4.5 ± 0.3 s^−1^, P710R 25 hep *k*_cat_ = 4.3 ± 0.3 s^−1^, p = 0.52) (**Figure 3C**). This suggests that P710R has a greatly reduced ability to access the sequestered folded-back state.

**Figure 3:**
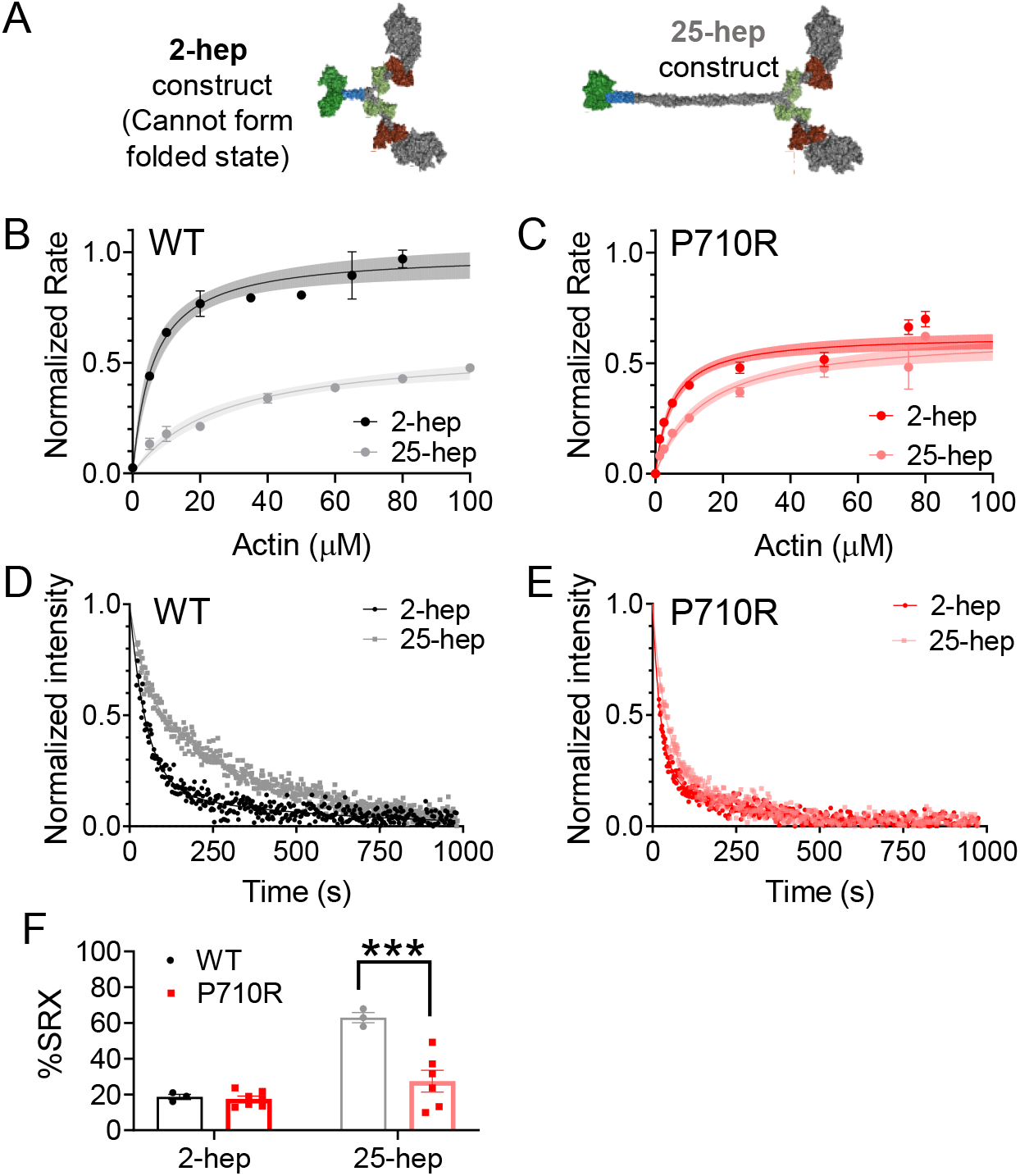
β-cardiac myosin ensembles with P710R mutation had reduced SRX state. **(A)** Schematic of protein constructs show 2 head (S1) domains (with light chains) and a 2 or 25-heptad tail domain. (**B-C**) Ensemble measurement of actin-activated ATPase rate normalized to the *k*_cat_ for WT 2-hep from preparations performed on the same day. See also methods and Figure S3. Error bars represent s.e.m. of measurements, and shading represents error of the fit. (**D-E**) Representative traces of mant-ATP single turnover in the short (2-hep) and long (25-hep) tailed protein constructs. (**F**) Quantification of the %SRX in WT and P710R myosins. Error bars represent s.e.m. WT data (D and F) was previously reported (15). *** indicates p < 0.001.

We have previously correlated the folded-back structural state with the proportion of basal ATP turnover at the SRX rate (~0.003 s^−1^), as opposed to the disordered relaxed state (DRX) rate (~0.01 – 0.03 s^−1^) (16, 53).These rates are measured using a single mant-ATP turnover assay in the absence of actin, in which the release of fluorescent mant-ADP from myosin is measured over time. The signal shows a biexponential decay characterized by two rates: the faster rate corresponds to the DRX rate, and the slower rate corresponds to the SRX rate (**Figure 3D**). Lower fractions of SRX correlate with a reduced ability to form the folded-back state (via EM imaging and ATPase (16)). We have previously showed that the percentage of WT 25-hep myosin hydrolyzing ATP at the slower SRX rate was 55-65% (at a salt concentration of 5 mM KOAc), in contrast to WT 2-hep myosin, which shows 15-25% SRX (regardless of salt concentration) due to its inability to form the folded-back state (**Figure 3F**) (15). Here, we found no significant difference in the percentage of molecules in the SRX state between WT and P710R 2-hep myosins (p = 0.12, Figure 3F). In contrast to WT 25-hep, only 27 ± 6% of the P710R 25-hep myosin hydrolyzed ATP at the slow rate (p = 0.0001 vs. WT 25-hep, **Figure 3F**). The fast rate of P710R 2-hep myosin was higher than that of WT 2-hep myosin (p = 0.004, Table S1), while the fast rate of P710R 25-hep myosin was not significantly different from that of WT 25-hep myosin. Consistent with our 25-hep ATPase findings, these single-turnover results confirm that the P710R mutation reduced myosin’s ability to form the folded-back SRX state.

To summarize our molecular studies, the P710R mutation significantly reduced the load sensitivity of the detachment rate from actin, step size, and unloaded motility, suggesting hypocontractility. However, the mutation also reduced the ability of myosin to form the SRX state, suggesting hypercontractility. Thus, to understand these competing effects, we next investigated the effects of this mutation in human cardiomyocytes.

### The P710R mutation increased traction force generation and cell size in micro-patterned hiPSC-CMs

We used CRISPR/Cas9 gene editing to insert a point mutation of proline to arginine in one allele of the MYH7 gene (Figure S4). We first measured the effects of the P710R mutation on cellular force generation using traction force microscopy (TFM). Isolated hiPSC-CMs at 35-40 days after cardiac differentiaton were cultured for 4-5 days on hydrogels of physiologic stiffness (10kPa) with matrigel bioprinted in a mature cardiomyocyte aspect ratio of 7:1 (32, 34). We have previously shown that this platform optimizes force generation and dramatically enhances hiPSC-CM maturation (32). Patterned hiPSC-CMs with the P710R mutation produced significantly higher total peak force than WT controls (**Figure 4 A-C**). Our single cell platform allowed assessment of >30 cells per group from three differentiation batches (**Figure 4C-F**) and provides spatially and temporally resolved measurements of traction force for each cell. Contraction duration – measured as the time between peak velocity of contraction and peak velocity of relaxation as previously described (34) – was significantly lengthened in the P710R cells (**Figure 4D**). P710R cells were significantly larger than the WT (**Figure 4E**), and the significant increase in force persisted after normalizing peak force to cell spread area (**Figure 4F**).

**Figure 4:**
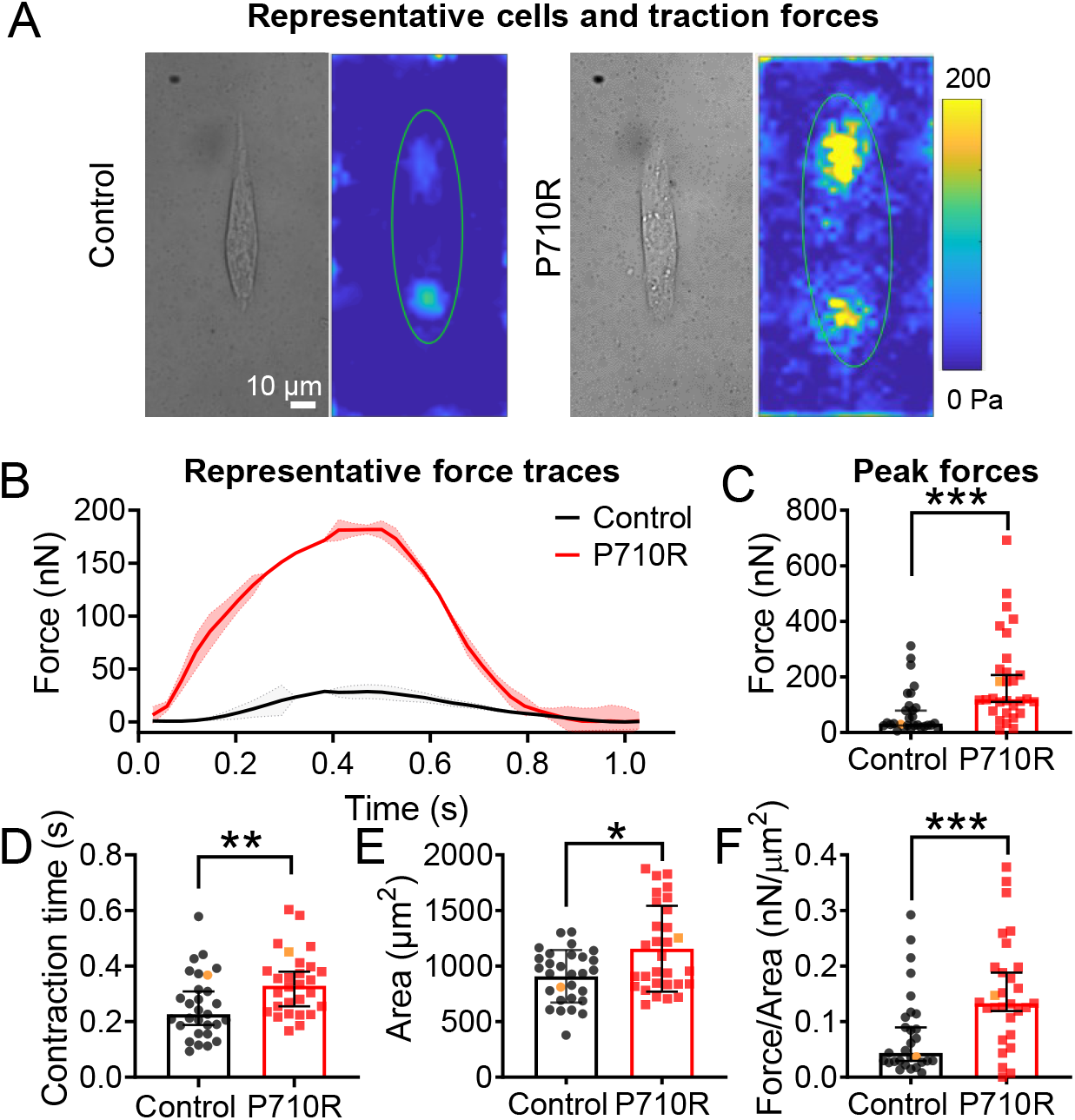
P710R mutation in micropatterned single hiPSC-cardiomyocytes significantly increased contractile function. (**A**) Representative single cells shown in bright field and as peak traction force plots. (**B**) Multiple (2-4) beats were captured per cell and the averaged traces for representative cells are shown, with shading representing the standard deviation of force between beats. (**C**) Peak total force and (**D**) contraction time of single control (n = 29) and P710R (n = 30) cells collected from 3 differentiation batches. (**E**) Cell spread areas were measured and used to calculate (**F**) cell force normalized to cell area. Representative cells are identified with orange markers in plots of population. Data are presented as median ± 95% confidence interval. * indicates p < 0.05, ** indicates p < 0.01, *** indicates p < 0.001.

### The P710R mutation caused myofibril disruption and z-disk thickening in patterned hiPSC-CMs

The ability of myosin to generate contractile forces *in vivo* depends on its incorporation into sarcomeres and myofibrils. Staining for β-MHC revealed organization into sarcomeres in cells grown on patterned hydrogels (**Figure 5A**). Sarcomere lengths were not significantly different between control and mutant cells when quantified from immunostained cells (**Figure 5B**). We confirmed this result using transmission electron microscopy (TEM) (**Figure 5C-D**). TEM of patterned cells showed aligned myofibrils in the WT controls and varying degrees of myofibril and sarcomeric disruption in many P710R cells. The degree of myofiber disarray was quantified using the directionality tool in ImageJ, and the degree of dispersion of directionality across the image was significantly increased in P710R cells (**Figure 5E**). We also observed significant thickening (measured along fiber direction) of the z-disks in P710R cells (**Figure 5F**). This cytoskeletal disruption may be an indication of cellular remodeling which ultimately leads to hypertrophy.

**Figure 5:**
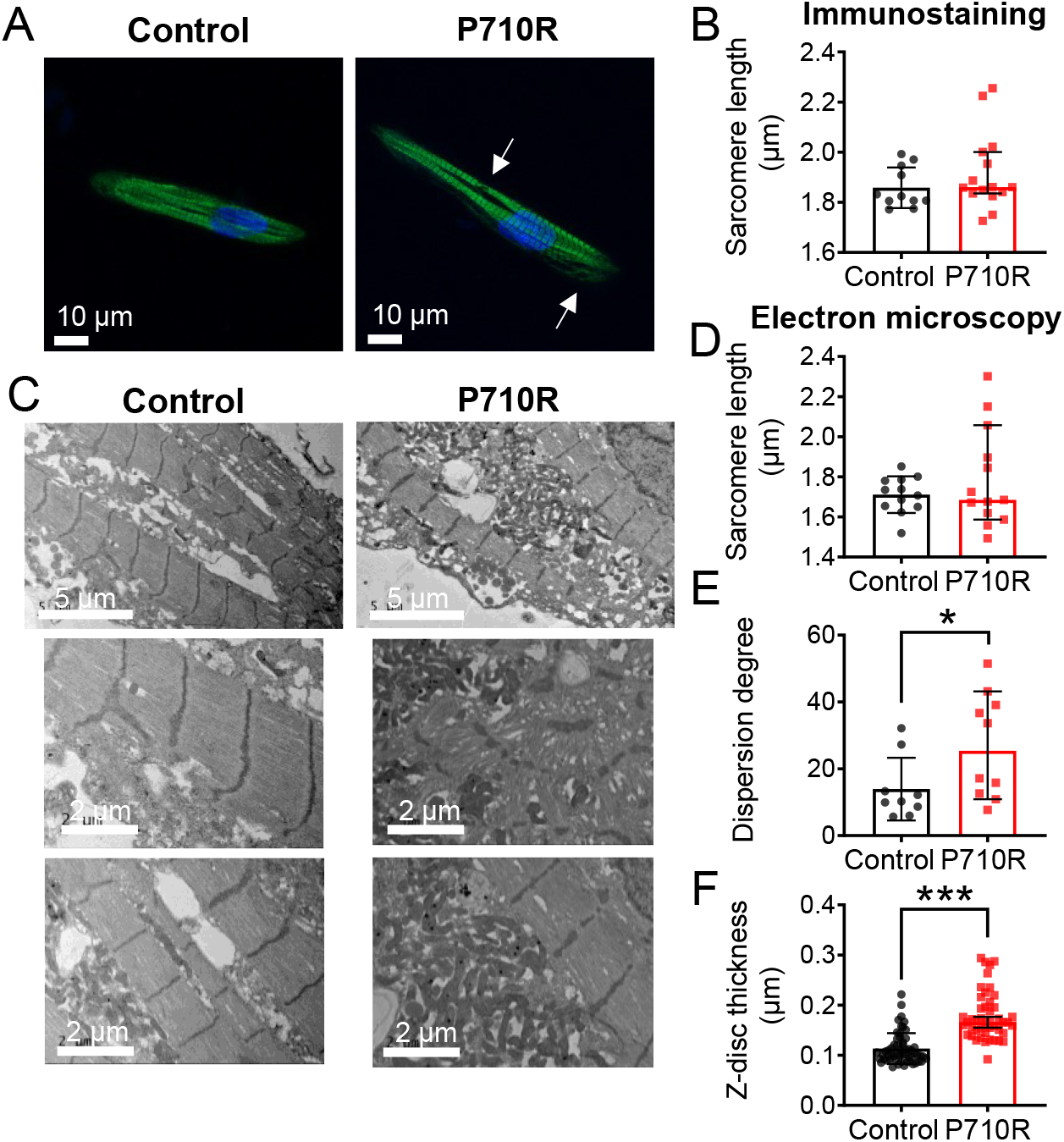
P710R mutation in micropatterned hiPSC-cardiomyocytes significantly disrupted myofibril organization. (**A**) Immunostaining of β-cardiac myosin in micropatterned cells (sarcomere disruption is marked with arrows) enabled (**B**) quantification of sarcomere length in micropatterned cells. (**C**) Representative transmission electron microscopy images of patterned cells enabled (**D**) quantification of sarcomere length, (**E**) dispersion degree, a measure of myofibril alignment, and (**F**) z-disk thickness from TEM images. Data are presented as median ± 95% confidence interval. * indicates p < 0.05.

### The P710R mutation caused hypertrophic growth of hiPSC-CMs via Akt and ERK

We next tested whether the P710R mutation increases cell size across the broader population of cells (and not only in paceable, patterned cells). After 45 days of culture in confluent monolayers, hiPSC-CMs were replated sparsely onto Matrigel coated tissue culture plastic, allowed to attach and spread for 2 days, and then fixed and stained to quantify their spread area. P710R cells were significantly larger than the control cells (**Figure 6A**). Western blot analysis of two known hypertrophic growth signaling pathways showed increased ERK and Akt basal activation (**Figure 6B-C**) in P710R cells. Inhibition of these signaling proteins with previously characterized specific inhibitors (SCH772984 and Akti 1/2) (54, 55) between day 26 and 46 resulted in significantly reduced cell area in P710R cells relative to untreated P710R cells (**Figure 6D**). However, the treatment of P710R cells with either inhibitor did not fully reduce their area to the size of the corresponding inhibitor treated unedited control cells (Figure S5).

**Figure 6:**
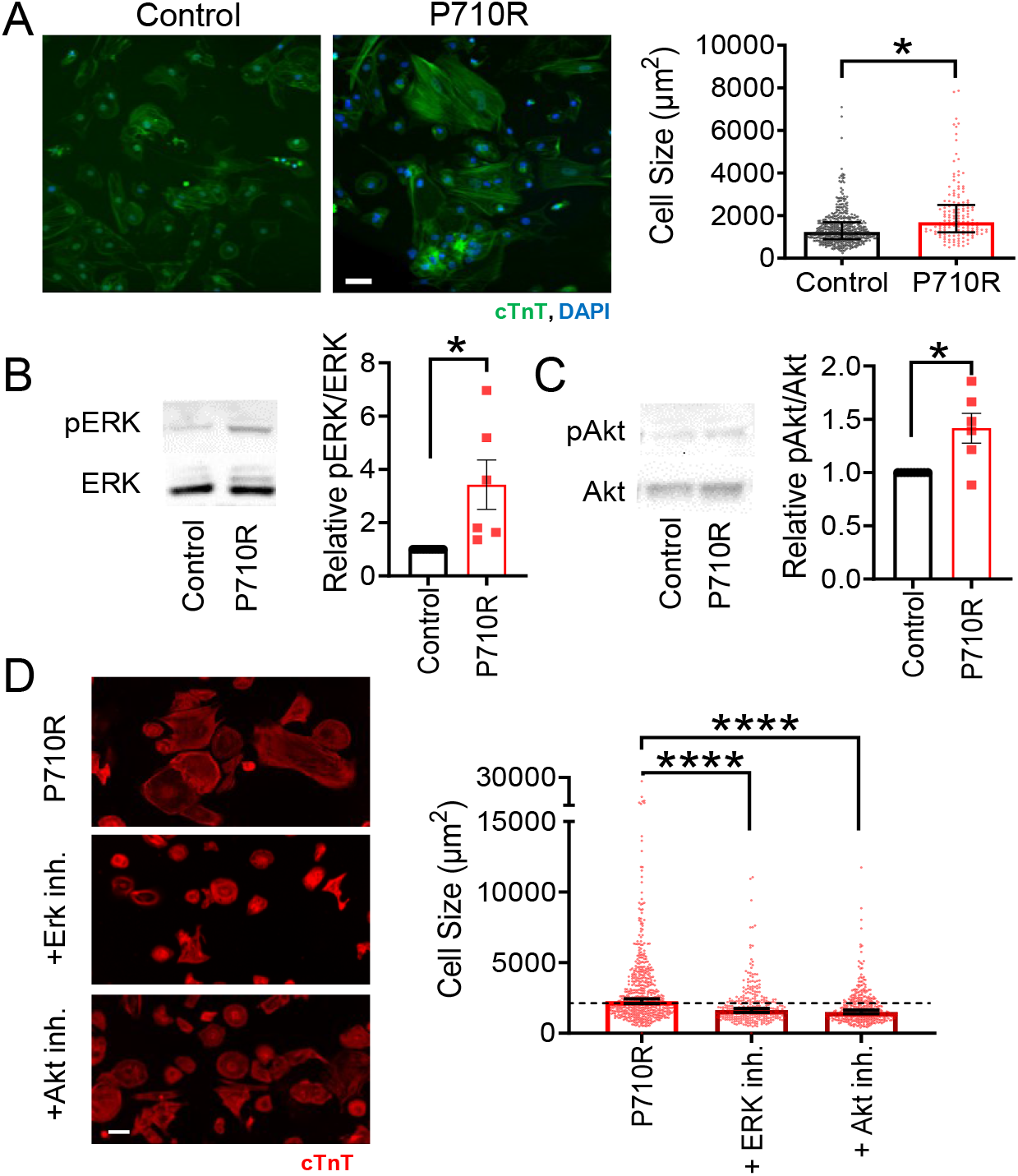
P710R mutation in hiPSC-cardiomyocytes significantly increases cell size and activation of proteins involved in hypertrophic signaling. (**A**) Size of unpatterned cells was quantified from immunostaining for cardiac troponin T (cTnT). (**B**-**C**) Western blots and densitometry analysis of phosphorylation of the hypertrophic signaling pathway proteins ERK (B) and (C) Akt in unpatterned cells with each P710R sample normalized to matched isogenic control. Data are presented as mean ± standard deviation. (**D**) Quantification of cell size after treatment of P710R cells with specific inhibitors of ERK and Akt. Scale bar represents 50 microns. Graphs of cell area are presented as median ± 95% confidence interval. * indicates p < 0.05 and *** indicates p < 0.001.

We also quantified the relative expression of β-MHC and α-MHC in P710R cells compared to isogenic control cells with both western blot and qPCR (Figure S6). We found no significant difference in β-MHC protein or transcript levels between P710R and control cells, while α-MHC protein and transcript levels were significantly reduced in P710R cells compared to control. However, the relative transcript levels of α-MHC to β-MHC was less than 0.1, suggesting that the majority of myosin heavy chain in both isogenic control and P710R cells was β-MHC.

### Computational modeling of P710R myosin compared to WT predicted an increase in force when SRX disruption was included

To determine the predicted effects of individual and combined changes in myosin kinetics on total force production, we used a previously validated computational model of cardiac muscle contraction (47), modified to contain an exponential load dependent detachment rate (**Figure 7A** and Figure S7). This model includes an OFF state to represent the SRX state of myosin. We measured calcium transients in monolayers of isogenic control and P710R cells when paced at 1 Hz (Figure S8), scaled the transients to the range of calcium concentration previously published for this model, and used the scaled transients as inputs for the calculations (56) (Figure S9A). We incorporated in this model the measured unloaded detachment rate (*k*_0_), its load sensitivity (*δ*), and step size (*x*_cb_). We defined the rate of myosin attachment (*k*_A_) from the previous modeling analysis that used our steady state actin-activated ATPase measurements for sS1 (26) (see methods for more details).

**Figure 7:**
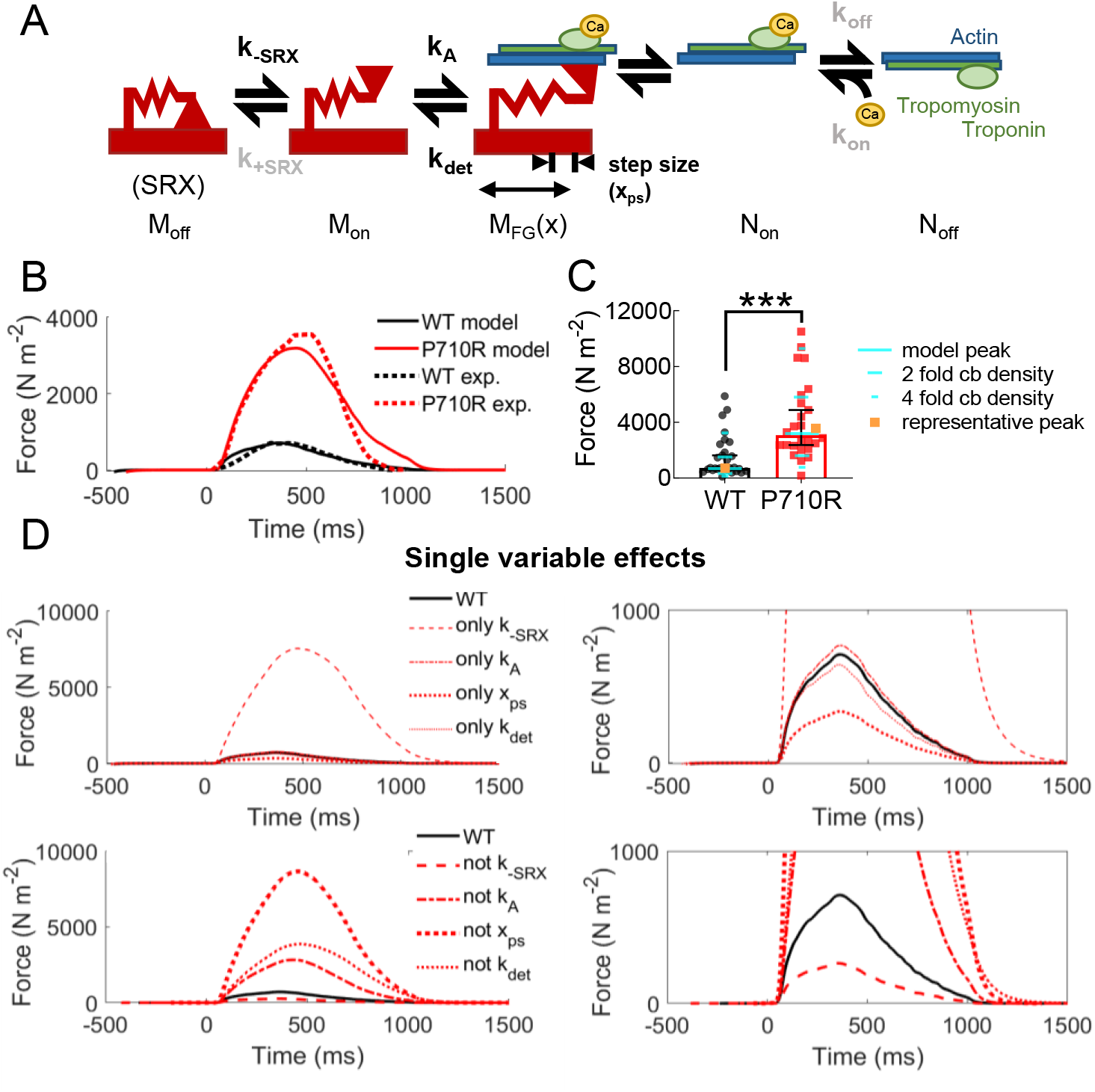
Modeling predicted that P710R induced modulation of SRX/DRX transitions are essential for hypercontractility. (**A**) Model schematic with adaptation of detachment rate to incorporate an exponential dependence on load. M_off_, M_on_ and M_FG_(x) represent the different states of myosin. N_on_ and N_off_ represent available and deactivated thin filaments, respectively. Parameters estimated or fit from in vitro experiments are shown in black, while rates held constant between groups are shown in gray. The model predicted (**B**) contractile force (solid line), which was fit to force traces from representative cells (dashed lines) normalized to cell cross-sectional area. (**C**) The predicted peak force (teal) fell close to the median measured force in our cell population when normalized to cross-sectional area, and two to four-fold changes to cross-bridge (cb) density relatively match the measured distribution of data. Representative cells highlighted in orange. (**D**) Simulations of independent effects of the four parameters (by inclusion with WT parameters and replacing from the best-fit mutant parameters). Zoomed in plots are included to improve visualization of low force traces.

We did not directly measure the rates of transitioning into and out of the SRX state, but we did measure significant differences in the percentage of myosins in the SRX state between WT and P710R in the presence or absence of actin (**Figure 3**). After identifying a set of parameters that successfully predicted the measured force output of a representative control cell (normalized to cross-sectional area), we found a best fit for the relative increase in rate of opening from the OFF state (*k*_−SRX_) to fit the force output of a representative P710R cell (see methods and Figure S9B). This process determined that a 12.9 fold (confidence limits 10.8-14.6 fold) increase in *k*_−SRX_ provided the optimal fit to the representative data. An increase in *k*_−SRX_ is consistent with the decrease in functional SRX state we measured experimentally, and the model predicted SRX% to be lower for the P710R cell at both contracted and relaxed periods of the cardiac cycle (Figure S9F). The rate of entering the SRX state (*k*_+SRX_) was set to 200 s^−1^ for both groups based on published data and previous modeling (47, 57). All other parameters were held constant between control and P710R, including the rates of thin filament activation (*k*_on_), inactivation (*k*_off_), and the cooperativity coefficient (*k*_coop_). The changes to myosin kinetics (*k*_−SRX_, *k*_0_, *δ* and step size) were sufficient to increase the percentage of thin filament activation and binding over the time course of contraction (Figure S9C).

The force predicted by the model after fitting parameters (Table S2) agreed well with the representative force traces (**Figure 7B**) and with the median force normalized to cross-sectional area of the population of cells (**Figure 7C**). Sensitivity analysis (Figure S10) suggested that four-fold changes in cross bridge density up and down spans the range of 90% of the cell forces we measured and matches the distribution of cell data. This parameter incorporates both the number of myosin heads per myofibril (which may vary with cell maturity and myofibril disruption) and the density of myofibrils in a cell (which we observed to be variable in these cells - Figure S11). Further simulations using the best fit parameter set and independently adding or removing each of the four measured parameters revealed that modulation of the rate of exiting the SRX state accounts for the majority of the measured increase in force (**Figure 7D**). An important note is that this analysis does not necessarily prove that the basal rate of opening from the SRX state is specifically increased, as opposed to an equivalent reduction in the rate of returning to the SRX state (Figure S9). Both our sensitivity analysis (Figure S10) and additional fitting suggest (Figure S9B) that the effects of changing *k*_−SRX_ and *k*_+SRX_ are nearly mirrored. Our results support the hypothesis that a change in the equilibrium between the SRX state and actively cycling states is essential for the P710R mutation to produce significant hypercontractility.

In summary, our cellular experiments confirm a hypercontractile and hypertrophic phenotype in hiPSC-CMs harboring the P710R mutation. Furthermore, our modeling results suggest that the measured molecular effects of P710R are consistent with hypercontractility if and only if SRX regulation is included.

## Discussion

The diversity of molecular alterations associated with HCM have made predictions of disease penetrance, severity, and response to therapeutics challenging. Our study demonstrates the power of an integrated, multiscale approach to study the biomechanical effects of mutations in myosin known to cause HCM. The P710R mutation was previously reported to have reduced ATPase activity and motility velocity (26), suggesting in isolation that it should reduce duty ratio and cause hypocontractility at the cellular level. In the present work, integrating molecular measurements of single molecules and ensembles of single and two-head constructs provides a more complete picture of the molecular mechanisms which contribute to the hypercontractility observed *in vivo*. By combining both a cellular model system and computational modeling, we were able to validate the hypercontractile effect of this mutation, demonstrate additional HCM-associated phenotypes, and provide key insights into the driving factors by which HCM mutations can result in hypercontractility.

Single molecules of β-cardiac myosin with the P710R mutation have reduced step size *d*, preserved actin detachment rate under zero load *k*_0_, and reduced load sensitivity *δ* of the actin detachment rate (**Figure 1**). This combination of findings suggests a possible molecular mechanism described as follows. In WT myosin, the converter domain, where P710 resides, couples the swing and forces at the lever arm to the conformational changes and forces at the nucleotide pocket. Through this coupling, load on the lever arm affects the rate of release of ADP from the pocket, which determines the actin detachment rate *k*_det_. In P710R myosin, the proline to arginine substitution may disrupt the local structure of the converter domain, resulting in a reduction in the described coupling. Consequently, the rate of ADP release (the actin detachment rate *k*_det_) is less dependent on load force (smaller *δ*) because load placed on the lever arm is no longer fully conveyed to the nucleotide pocket. Similarly, the step size *d* is reduced because the conformational change at the pocket is no longer fully conveyed to the lever arm. The unloaded ADP release rate (unloaded actin detachment rate *k*_0_) is not affected by the mutation presumably because the local structure of the nucleotide pocket remains intact. Due to inherent variability among individual molecules, as evident in our single molecule data (**Figure 1**), the extent of disruption by P710R may vary. Thus, P710R may cause a small percentage of molecules to be fully dysfunctional in moving actin (“deadheads”), resulting in the observed higher numbers of immobile actin filaments in the motility assay (Figure S2; see also methods).

We have previously observed reductions in the load sensitivity of myosin’s actin detachment rate by cardiac myosin effectors but not by cardiomyopathy causing mutations (12). For example, the investigational heart failure drug omecamtiv mecarbil (OM) dramatically reduces the detachment rate, its load sensitivity (12), and myosin’s step size (50). As the effects of OM and P710R share some similarities, it is important to note that residue P710 is part of the OM binding pocket in the converter domain (58). A recent preprint reported a reduction in myosin’s step size due to a different HCM mutation in the same region (P712L) (59). Taken together, these results emphasize the converter domain’s critical facilitation of myosin’s powerstroke and load sensitivity and reveal various consequences of perturbing this domain.

A major finding of this study is that the disruption of the folded, SRX state is a critical driver of hypercontractility with the P710R mutation. The results of the actin-activated ATPase and single-turnover experiments using the longer tailed 25-hep construct (**Figure 3**) demonstrate that a significantly larger fraction of myosins with this mutation are in an open, more active state than WT control. This increases the number of heads available for binding to actin, which we hypothesized would increase force production. Interestingly, this mutation lies neither in the myosin mesa region nor the predicted head-head interaction site of existing folded-back myosin structural models (18, 60–62). This suggests that while it may not directly impact interactions required to form the folded state, the P710R mutation may result in allosteric changes that indirectly reduce myosin’s ability to form the folded SRX state. We have measured similar changes in SRX across a range of HCM-causing mutations (15, 17), and others have confirmed these effects in cellular and animal models (16, 63), suggesting that SRX regulation may be a major cause of hypercontractility in HCM. This is further corroborated by the finding that mavacamten, a small-molecule inhibitor of myosin that is known to increase the SRX state, is effective in reducing symptoms in patients with HCM (16, 64). Our computational modeling analysis also supports the importance of destabilization of the SRX for increasing net force even when combined with a hypocontractile change like decreased step size.

We confirmed that the P710R mutation causes hypercontractility and other key features of HCM in cardiomyocytes using a cellular experimental model in parallel with molecular level experiments. Using a micropatterned culture system also allowed for measurement of cellular scale forces that are enhanced by the alignment of myofibrils and balanced by a physiologic-range substrate stiffness (10 kPa). This system has also been used to quantify hypercontractility in hiPSC-CMs with HCM-causing troponin mutations (65). The force per cross-sectional area we measured in the cells was significantly lower than what has been reported for mature cardiac tissue, but this is a known sign of the relative immaturity of hiPSC derived CMs (66). Despite this limitation, cells with the P710R mutation produce significantly greater force even after normalizing for the increase in cell area (**Figure 4**). There was significant variability in force production within the cell populations, which may be due in part to differences in myofibril density between cells, as suggested by our modeling analysis. The P710R mutation caused significant perturbation in alignment of myofibrils in the mutant cells, a well described phenotype in HCM (67), as well as an increase in the thickness of the z-disc (**Figure 5**). Increased z-disc thickness has been described in many cardiomyopathies (68–70), which suggests that z-disc thickening is a conserved phenotype associated with hypercontractility and hypertrophic remodeling. Finally, we observed cellular hypertrophy which was mediated in part by Akt and ERK. ERK activation has been specifically linked to concentric hypertrophy in response to an increase in force index (force production integrated over time of contraction) (71). By comparing cellular responses to an isogenic control, we have more confidence that the altered biomechanical consequences of the P710R mutation are driving both hypercontractility and additional cellular dysfunction.

The P710R mutation induces a number of profound changes to myosin activity whose effects become easier to interpret when integrated into a computational model. Even with similar driving calcium transients and identical rates of thin filament activation by calcium, the model simulations predict an increase in thin filament activation and a lengthening of contraction time resulting solely from the changes in myosin kinetics (specifically the change in regulation of SRX). In simulations where only step size, attachment and detachment rates were changed (as is true for the sS1 construct), there were minimal effects on actin availability (±15% change), which matches the observed similarity in pCa_50_ in sS1 motility with or without the mutation. When the disruption of SRX caused by the mutation is included, the predicted percentage of actin kept in the active state by myosin increases by 8-10 fold. The model used in this study, while by no means comprehensive, provides valuable insights into the expected individual and combined effects of changes to myosin kinetics and biomechanical function on force production, actin activation, and potential sources of variability.

Several important limitations of this project should be addressed. We measured the SRX state using shortened constructs that do not contain the full thick filament backbone and thus likely underestimate the additional stabilization of the SRX in thick filaments in vivo. Hypertrophy measurements and assessment of protein activation were performed on unpatterned cells grown on tissue culture plastic or glass because it would be difficult to quantify phosphorylation in small populations of cells recovered from patterned gel platforms. This modeling work shows the singular effects of a mutation, but does not fully capture potential synergistic or emergent consequences of a mixture of control and mutant myosins acting together. Furthermore, this modeling approach assumes that muscle force is felt uniformly throughout the muscle, while in reality, the spatial distribution of myosins within a sarcomere can have profound effects on the force experienced by individual myosin heads. Future modeling could incorporate spatially explicit information that could provide even more detail on the effects of local loads and heterotypic myosin effects.

In summary, the P710R mutation reduces several functional parameters of the isolated myosin molecule; however, the dysregulation of the SRX state appears to be the major driver of hypercontractility. When expressed as a heterozygous mutation in human stem cell derived cardiomyocytes, this mutation causes hypercontractility, myofibril disruption, and hypertrophy mediated in part through ERK and Akt. This project demonstrates the value of computational modeling integrated with multiscale measurements to give insights into the interactions of load-sensitive, functional parameters and provides a platform to study other mutations and therapeutics.

## Materials and Methods

### Molecular measurements of myosin function in transgenic human proteins with P710R mutation

Recombinant human β-cardiac myosin protein constructs (short subfragment 1 (sS1), short-tailed (2-hep), and long-tailed (25-hep)) were expressed in C2C12 mouse myoblast cells and purified as previously described (11, 18). The load-dependent detachment rates of WT and P710R sS1-eGFP molecules were measured in a dual-beam optical trap using the harmonic force spectroscopy (HFS) method previously described (12, 48, 72) with slight modifications. The step sizes of myosins were determined from the same HFS data by adapting the ensemble averaging method (73) to HFS’s oscillatory data. For each molecule, position traces of all events were aligned at the start of binding, extended to the length of the longest event, and averaged. Then the oscillations were removed by subtracting a fitted sine function. The total step size for each molecule was taken as the difference between the initial position and the end position of the extended, averaged, sine-subtracted traces. Motility measurements of WT and P710R sS1-AC were performed using our previously described motility assay (29, 74) with some modifications. Motility measurements with regulated thin filaments were performed using recombinant human troponin complex and bovine cardiac tropomyosin (75). NADH-coupled ATPases were used to compare the activity of the 2-hep and 25-hep constructs of WT and P710R β-cardiac myosin as previously described (76). Single turnover experiments were performed as previously described (15, 16), using WT and mutant versions of human ß-cardiac 2-hep and 25-hep. Additional details are included in the data supplement.

### Measurements of functional consequences of P710R mutation in gene edited hiPSC-CMs

The control human iPSC line (Stanford Cardiovascular Institute [SCVI-113]) was obtained from the Stanford CVI iPSC Biobank. The P710R mutation was edited into these cells using a method that has been described previously (77, 78). Cells were differentiated and their contractile function was assessed on micropatterned hydrogels gels as previously described (32, 34). Cells on micropatterened gels were fixed and stained for β-cardiac myosin and their sarcomere length quantifed. Cells were also plated onto micropatterned film and imaged with transmission electron microscopy. Unpatterned cells were stained for cardiac troponin T to measure cell area. To clarify the effect of ERK and Akt, the cells were treated with either Akt inhibitor (Akti 1/2, Tocris: 5μM) or and ERK inhibitor (SCH772984, SelleckChem: 1μM) every 3 days between day 26 and day 46 (a time frame over which hiPSC-CMs have been shown to increase in area). Additional details are included in the data supplement.

### Implementation of computational model

We used a previously validated continuum model which incorporates discretized flux equations to describe transitions of myosin heads and thin filaments from inactive to active/bound states and provides prediction of active and passive force (47, 79). The rate of myosin detachment from actin was adapted to match the measured exponential load dependence, and the relative change in k-SRX was fit using experimental force measurements from representative hiPSC-CMs. Additional details about model implementation are included in the data supplement.

## Acknowledgements

This work was funded by NIH grants RM1GM33289 to JAS, DB, and BLP; 1R21HL13099301 to DB and BLP; HL117138 and S10RR026775 to JAS; HL123655 to DB; and HL133359 and HL149164 to KSC. ASVR and KBK were supported by NIH T32 HL094274. It was also funded by American Heart Association grants 17CSA33590101 to BLP and DB; and American Heart Association Postdoctoral Fellowship (20POST35211011) to ASVR. MMM was supported by the Stanford Cellular and Molecular Biology training grant. GP was supported by the Swiss National Science Foundation (SNSF) Early Postdoc Mobility Fellowship (#P2SKP2_164954).. The human iPSC line(s) were obtained from Joseph C. Wu, MD, PhD at the Stanford Cardiovascular Institute funded by NIH 75N92020D00019. We acknowledge John Perrino and the electron microscopy imaging core for their help with fixing, sectioning and imaging of the cell samples; Alexandre Ribeiro for his initial help with micropatterning experiments; Arjun Adhikari for his preliminary work on the molecular data; a generous gift of an α myosin antibody from Theresia Kraft; and Oscar Abilez for his help with imaging calcium transients.

