## Supplementary Information for "Hypertrophic cardiomyopathy ß-cardiac myosin mutation (P710R) leads to hypercontractility by disrupting super-relaxed state"

**This PDF file includes:**

Supplementary text
Figures S1 to S11
Tables S1 to S3
Legends for Movies S1 to S2
SI References

**Other supplementary materials for this manuscript include the following:**

Movies S1 to S2

### Detailed Methods

#### *Expression and purification of proteins*

Recombinant human β-cardiac myosin protein constructs – short subfragment 1 (sS1), short-tailed (2-hep), and long-tailed (25-hep) – were coexpressed with a FLAG-tagged human essential light chain (ELC) in mouse myoblast cells (C2C12) using adenoviral vectors. The sS1 constructs had either a C-terminal enhanced green fluorescent protein ("sS1-eGFP") or a C-terminal eight-residue PDZ-binding peptide ("sS1-AC"). The sS1 constructs were purified with a FLAG affinity and ion-exchange chromatography as previously described (1). The 2-hep and 25-hep constructs had both an eGFP and PDZ tag on their C-termini. For these longer constructs, human RLC was exchanged for the endogenous mouse RLC while bound to a FLAG resin, and then the myosin was purified as previously described (2).

Actin was prepared as described previously (3) from bovine cardiac muscle (the sequences are identical between bovine and human actin). The actin was dialyzed in ATPase buffer (5 mM KCl, 1 mM DTT, 3 mM $MgCl_2$ and 10 mM imidazole) to form F-Actin for ATPase assays.

#### *Deadheading myosin*

Myosin protein used for unloaded motility experiments was further subjected to a 'deadheading' procedure to remove inactive heads. Myosin was mixed with 10x excess of unlabeled F-actin on ice for 5 min, followed by addition of 2 mM ATP for 3 min, then centrifuged at 95 krpm in a TLA-100 rotor (Beckman Coulter) at 4 °C for 20 min. The supernatant was collected. We found that the P710R protein had more inactive heads than WT, as seen in a greater percentage of stuck filaments (5-15%) even after deadheading two times (Figure S2). In contrast, percent stuck filament was 0-5% for WT sS1-AC with or without deadheading (Figure S2). Note, however, that these stuck filament percentages do not directly translate to the percentage of inactive heads; only a very small number of inactive heads (probably 1% or less) is needed to produce a much larger number of stuck filaments. For RTF motility experiments, we skipped deadheading because it did not substantially reduce percent stuck filaments, as 5-15% remained. This is likely because deadheading cannot remove 100% of completely or partially inactive heads.

#### *Single-molecule measurements of load-dependent detachment rates and step sizes with dual-beam optical trap*

The load-dependent detachment rates of WT and P710R sS1-eGFP molecules were measured in a dual-beam optical trap using the harmonic force spectroscopy (HFS) method previously described (4-6) with slight modifications. The sample chamber consisted of two double-sided tapes between a glass slide and a coverslip spin-coated with 1.6-um-diameter silica beads (Bang Laboratories) and a solution of 0.1% nitrocellulose/0.1% collodion in amyl acetate. Experiments were done at 23 °C. The following solutions were flowed into the chamber sequentially, each with a couple minutes of incubation: 1. GFP antibody (Abcam ab1218) diluted to 1 nM in assay buffer (AB; 25 mM imidazole pH 7.5, 25 mM KCl, 4 mM $MgCl_2$, 1 mM EGTA, and 10 mM DTT). 2. BSA diluted to 1 mg/mL in AB (ABBSA), to wash and block the surface. 3. Human β-cardiac myosin sS1-eGFP diluted to 10-50 nM in ABBSA, to bind all the GFP antibodies on the surface. 4. ABBSA, to wash. 5. A final solution consisting of 2 mM ATP, 0.3 nM TMR-phalloidin-labeled biotinylated actin (Cytoskeleton) filaments, an oxygen-scavenging system (0.4% glucose, 0.11 mg/mL glucose oxidase, and 0.018 mg/mL catalase), and 1-um-diameter NeutrAvidin-coated polystyrene beads (ThermoFisher) or 1-um-diameter streptavidin-coated polystyrene beads (Bang Laboratories) diluted in ABBSA. The chamber was sealed with vacuum grease and used for up to 2 hr.

The two traps were calibrated using the power spectrum method with corrections for filters, aliasing, and surface effects (7), and their stiffnesses were 0.08-0.10 pN/nm in these experiments. After calibration, a "dumbbell" was made by stretching an actin filament between the two trapped beads (Figure S1A). While the stage oscillated at 200 Hz, the dumbbell was lowered near platform beads on the surface to test for binding to a potential myosin. Robust interactions with myosin were indicated by consistent, large, brief displacements of the trapped beads due to stronger association with the oscillating stage. We typically explored ~4-10 platform beads before robust interactions were observed, suggesting that the GFP antibody concentration used resulted in sufficiently low numbers of properly oriented molecules on the surface for single molecule conditions. Time traces were saved and automatically processed as previously described (4, 6). Briefly, events were selected based on a simultaneous increase in oscillation amplitude and decrease in phase between each bead and the stage (Figure S1B). The detachment rate at each force was determined by a maximum likelihood estimation from the durations of events within the force bin. The rates at different forces were then fitted to Eqn. 1 to obtain the two parameters $k_0$ and $\delta$. The data on multiple molecules presented in this paper are from multiple dumbbells in independent experiments from multiple days. Statistical significance of the difference in the weighted mean of these parameters between WT and P710R was determined using a two-tailed unequal variances t-test.

The step sizes of myosins were determined from the same HFS data by adapting the ensemble averaging method (8) to HFS's oscillatory data. For each molecule, position traces of all events were aligned at the start of binding, extended to the length of the longest event, and averaged (Figure S1C and D). Then the oscillations were removed by subtracting a fitted sine function (Figure S1D). The total step size for each molecule was taken as the difference between the initial position and the end position of the extended, averaged, sine-subtracted traces. Statistical significance of the difference in step size between WT and P710R was determined using a two-tailed unequal variances t-test.

#### *In vitro motility assays*

Motility measurements of WT and P710R sS1-AC used our previously described motility assay (9, 10) with some modifications. Multi-channel flow chambers consisted of double-sided tapes between a glass slide and a coverslip pre-coated with a solution of 0.1% nitrocellulose/0.1% collodion in amyl acetate.

For unloaded motility, the following solutions were flowed into the chamber sequentially: 1. SNAP-PDZ18 (purified as described in Aksel et al. *Cell Reports* 2015 (9)) diluted to 3 μM in assay buffer (AB; 25 mM imidazole pH 7.5, 25 mM KCl, 4 mM $MgCl_2$, 1 mM EGTA, and 10 mM DTT), incubated for 2 min. 2. BSA diluted to 1 mg/mL in AB (ABBSA), incubated for 2 min. 3. Human β-cardiac sS1-AC diluted to ~200 nM in ABBSA, flowed twice with 2 min incubation each time. 4. ABBSA. 5. A final solution consisting of 2 mM ATP, 2.5 nM TMR-phalloidin-labeled actin filaments, and an oxygen-scavenging system (0.4% glucose, 0.11 mg/mL glucose oxidase, and 0.018 mg/mL catalase) in ABBSA. pH of the final solution was measured to be 7.5 at 23 °C. See Movies S1 and S2 for example motility measurements of WT and P710R, respectively.

For RTF motility, a full pCa curve was generated by one slide with 6-7 channels. The following solutions were flowed into the channels sequentially: 1. SNAP-PDZ18 diluted to 3 μM in AB, incubated for 2 min. 2. ABBSA, incubated for 2 min. 3. Human β-cardiac sS1-AC diluted to 400-600 nM in ABBSA, flowed twice with 2 min. incubation each time. 4. ABBSA. 5. TMR-phalloidin-labeled actin filaments diluted to 10 nM in ABBSA, incubated for 1 min. 6. ABBSA. 7. A final solution consisting of 2 mM ATP, an oxygen-scavenging system (0.4% glucose, 0.11 mg/mL glucose oxidase, and 0.018 mg/mL catalase), 100 nM human troponin complex (expressed, purified, and complex formed as described in Sommese et al. *PloS One* 2013 (11)), 100 nM bovine cardiac tropomyosin (purified as described in Sommese et al. *PloS One* 2013 (11)), and varying amounts of $CaCl_2$ (as calculated in (12)) in ABBSA. pH of the final solution was measured to be 7.5 at 23 °C. Troponin and tropomyosin were added only in the final solution because we found that filaments were decorated with the regulatory proteins very quickly and certainly by the time the channels were imaged.

The slide was then placed on a Nikon Ti-E inverted microscope with a 100x TIRF objective. Images were taken for 30 sec at 2 Hz with 300 ms exposure on an Andor iXon+EMCCD camera. Three movies were recorded for each channel. The objective temperature was maintained at 23 +- 0.3 °C as closely as possible to minimize variations in velocities stemming from myosin's high sensitivity to temperature. Movies of all channels on each slide were acquired as quickly as possible, within ~15 min after flowing in the final buffer for the 6-7 channel slides. Despite our foregoing an ATP regeneration system, no slowing down of actin filaments was observed throughout imaging, nor any differences observed with or without an ATP regeneration system.

Movies were processed by FAST (Fast Automated Spud Trekker) for filament tracking and velocity analysis (9) (Figure S2A and B). The following parameters were used in FAST: window size n = 5, path length p = 10, percent tolerance pt = 80, and minimum velocity for stuck classification minv = 20 nm/s. All velocities reported in the main text are the mean velocities including stuck filaments (MVIS), which is equivalent, up to a multiplicative factor, to the "percent time mobile" metric defined previously (9). The lower minv value and higher pt value as used here are appropriate for analyzing loaded or RTF motility data. While a higher minv (80, as used in some previous publications) and lower pt (20 in some previous publications) are more appropriate for analysis of unloaded motility data (along with reporting "Top5%" (Figure S2C) or "MVEL-filtered" velocities), we used the same set of FAST parameters to analyze all motility data reported in this paper so that values can be directly compared.

For RTF motility, the Hill equation was fitted for each independent experiment producing MVIS values at 6-7 pCa concentrations (6-7 channels). Variation among the three movies taken for any particular channel was very minimal and smaller than variation between independent slides/experiments. Therefore, to capture the true experimental or biological variation, 4-5 independent experiments (slides) were performed for both WT and P710R. Error bars on data points at each pCa value presented in Figure 2C represent s.e.m. of all independent experiments. Similarly, error bars on the fitted $pCa_{50}$ and $n$ values reported in Table 1 represent s.e.m. of the various fitted values from independent experiments.

Statistical significances of the differences in velocities or fitted values between WT and P710R were determined using two-tailed unequal variances t-tests.

#### *Actin-activated ATPase assay*

NADH-coupled ATPases were used to compare the activity of the 2-hep and 25-hep constructs of WT and P710R β-cardiac myosin as previously described (13). Briefly, actin was prepared as described in (3) and dialyzed 4x into ATPase reaction buffer without ATP: 5 mM KCl, 10 mM imidazole pH 7.5, 3 mM $MgCl_2$, and 1 mM DTT. Gelsolin was prepared as previously described (14) and added to the actin at a final molar ratio of ~1:200. In a 96-well plate, actin was mixed with ATPase reaction buffer to achieve final concentrations of 1-100 µM, and myosin was added at a final concentration of 0.025 µM. To measure the basal rate without actin, myosin was added to ATPase reaction buffer at final concentrations of 0.075 – 0.125 µM. To initiate the reaction, a 5x coupling solution containing 100 U/mL lactate dehydrogenase (Sigma L1254), 500 U/mL pyruvate kinase (Lee Biosolutions 500-20), 2.5 mM phospho(enol) pyruvate (Sigma P0564), 10 mM ATP, and 2 mM NADH (Sigma N8129) was added at 1/5 the volume of the final reaction. The plate was shaken for 5 minutes at 23°C before reading at 340 nm every 30 seconds for 25 minutes. A standard curve of ADP from 0 – 300 μM was created to convert the absorbance values to concentration of ADP produced.

Data for P710R was collected from triplicate measurements of two independent protein preparations, where both P710R 2- and 25-hep were prepared in parallel with WT 2-hep as a control. Each independent protein preparation was fit to the Michaelis-Menten equation using the non-linear fit feature in PRISM, allowing for calculation of the maximum actin-activated ATPase activity rate ($k_{cat}$) and Michaelis-Menten constant ($K_M$) (Table S1 and Figure S3). To determine data reported in Table S1, we averaged these values from the two independent protein preparations, weighting based on the standard error of the fit. To determine p-values, a paired t-test was used. A representative example of this raw data is shown in Figure S3, where error bars represent the standard deviation of the triplicate measurements and the shaded area represents the 95% confidence interval. To generate the normalized plot shown in Figure 3C, each independent protein preparation was normalized to the $k_{cat}$ of its same-day WT 2-hep control, then data from these two protein preparations was combined and plotted on the same graph. Error bars represent standard deviation, and the shaded area represents the 95% confidence interval. WT 2-hep is not shown on this plot for clarity. By contrast, the WT 25-hep data (Fig. 3B) represents triplicate measurements from a single protein preparation, since this data only aims to replicate our previous results using this NADH-coupled ATPase. Errors on the WT 25-hep data in table S1 represent standard error of the fit from a single measurement. Figure 3B shows the WT 25-hep data plotted with and normalized the $k_{cat}$ of its same-day WT 2-hep control.

#### *Mant-ATP single turnover assay*

Single turnover experiments were performed as previously described (15, 16), using WT and mutant versions of human ß-cardiac 2-hep and 25-hep. These experiments were performed in a 96-well plate by mixing 200 nM myosin in a buffer (10 mM Tris pH 7.5, 4 mM $MgCl_2$, 1 mM EDTA, 1 mM DTT, 5 mM potassium acetate) with 2′-(or-3′)-*O*-(*N*-Methylanthraniloyl) adenosine 5′-triphosphate (mant-ATP, Thermo Fischer Scientific) to a final concentration of 200-240 nM. ~10 s after mixing myosin and mantATP buffer, 2 mM ATP was added and the fluorescent signal (470 nm Em/405 nm Ex) was measured every ~2 s for 16 min. The traces were normalized, plotted, and fit to a five parameter bi-exponential decay function to yield rates and amplitudes of fast (DRX) and slow (SRX) phases (Table S1). Statistical significance was determined by two-way ANOVA and specific differences assessed with Sidak's multiple comparisons test.

#### *CRISPR/Cas9 based editing of hiPSCs*

The control human iPSC line (Stanford Cardiovascular Institute [SCVI-113]) was obtained from the Stanford CVI iPSC Biobank. These cells were acquired, reprogrammed and deidentified in accordance with approved IRB protocols and cultured in the appropriate media to maintain stemness. The iPSCs were maintained in defined E8 media (Gibco) on tissue culture plates coated with Matrigel (Corning) diluted 1:100 in DMEM/F12 at 37°C in 5% $CO_2$. The P710R mutation was edited into these cells using a method that has been described previously (17, 18). We designed a guide RNA (GCTGCGCTGCAATGGTGTGCTGG) that was predicted to be highly specific to a site 34 basepairs upstream of the desired mutation site and cloned it into the pSpCas9 (BB) 2A-GFP (PX458) plasmid. We also designed 500 basepair homology regions surrounding our cut site and including a C->G mutation at basepair 9699 to encode the P->R transition and a silent mutation to form a TTAA site at 9647-9650. These homology regions were cloned into an MV-Puro-TK plasmid surrounding a Puromycin resistance – TK selection cassette and both plasmids were introduced into the control iPSCs using Lipofectamine 3000. Puromycin was used to select for positively edited cells and then the Transposagen X reagent was used to excise the cassette leaving "scarless" silent TTAA mutation and Ganciclovir was used to select for successfully repaired cells. Sequencing confirmed correct, single allele edits with no indels or other mutations on the second allele of MYH7 or on MYH6 (a close homologue in this region).

#### *Cardiomyocyte differentiation and culturing of hiPSCs*

Cells were differentiated into the cardiac lineage following a well-established protocol that relies on sequential activation and inhibition of the Wnt pathway. Specifically, we began differentiation when the cells reached ~75% confluence in RPMI media with B27 minus insulin supplement and 5 μM Chir-99021. After 24 hours, the media was replaced or supplemented with RPMI media with B27 minus insulin supplement, and then at 48 hours, the media was replaced by RPMI media with B27 minus insulin supplement plus 5 μM IWR or 2 μM C59. On the fourth day after the start of differentiation, the media was replaced with RPMI media with B27 minus insulin supplement, and on the sixth day, the media was replaced with RPMI media with B27. Glucose starvation in glucose free RPMI with B27 was performed after the onset of beating for 4 days, and cells were replated before day 20 to allow for firm attachment to a fresh Matrigel coated plate. Cells were maintained in RPMI media with B27 for up to 60 days. Calcium transients were captured using a wide-field fluorescent camera on cells loaded with Fluo4 dye after 30 minutes using a point electrode and 1Hz pacing. Linear regions of interest approximately the length of 2-5 cells were chosen perpendicular to the direction of conduction at 4 different sites at least 2 mm apart. Transients were normalized to their baseline and the time to 90% of peak, time over 90%, and decay from 90% of peak to 5% above baseline times were measured and found to be not significantly different between control and mutant cells (Figure S8).

#### *Micropatterning hydrogels and substrates for microscopy*

Micropatterned hydrogels gels were created as previously described (19, 20). SU-8 photolithography was used to create molds for stamps with rectangular features 10 μm deep with a 7:1 aspect ratio and an area of 2500 $\mu m^2$. These single cell patterns were arranged in a 1 cm by 1 cm grid, spaced to minimize the risk of mechanical interaction between adjacent cells. The area of these patterns was also larger than the median and 75 quartile of the cell spread area for both control and P710R cells, so it allowed for assessment of cell area with a spatial cue. Polydimethylsiloxane (PDMS) was mixed at a 9:1 by weight ratio of elastomeric base to primer and curing agent, mixed, degassed, and cured at 65°C for an hour. The PDMS stamps were coated in a 10:1 dilution of Matrigel in L15 media overnight at 4°C, rinsed in media and gently dried with Nitrogen. Stamps were then inverted and pressed onto cleaned and plasma-treated glass coverslips or Aclar film and weighted with a 50 g mass for at least 3 minutes to ensure even pressure and coverage.

For creation of micropatterned hydrogels, a mixture of Acrylamide, Bisacrylamide, HEPES and 0.5 μm fluorescent beads (Life Technologies, FluoSpheres) when used for traction force microscopy and polymerized with TEMED and 10% APS according to a formula which has been validated to produce gels of a stiffness of 10 kPa. This stiffness matches the physiologic stiffness of healthy myocardium near the beginning stages of disease.

#### *Traction force microscopy and calculation*

Single cell traction forces were assessed from single cells using algorithms described in our previous publications. We used both a Leica live cell imaging scope with a Prime 95 camera and a Nikon TiE microscope with an Andor Neo sCMOS camera. Cells were imaged in the bright-field to determine cell area and screen for beating, apparently single cells. Videos were then collected in the bright field and fluorescent channels to capture at least two full cardiac contraction cycles, at a minimum of 25 frames per second using a 40x 0.85NA or 20x air objective.

Electrical field stimulation of 10V/cm at a frequency of 1 Hz was imposed with a carbon electrode throughout the acquisition of videos, and only cells that were able to keep up with 1 Hz pacing were included in further analysis. Our custom script and user interface in MATLAB allows for measurement of the cell outline and outputs the normalized displacements, total traction force, contraction time, moments, and integrated contraction impulse (force integrated over the time of contraction). The peak forces from these cells were not normally or log-normally distributed (according to Anderson-Darling, D'Agostino & Pearson, and Shapiro-Wilk tests), so statistical significance for these parameters was determined with non-parametric Mann-Whitney tests.

#### *Immunostaining and electron microscopy*

At day 45 after differentiation, the cells were replated on 48-well tissue culture plastic, hydrogels or micropatterned aclar film and allowed to attach and grow for 4 or 7 days before fixing for further imaging. For immunostaining to determine cell size, cells were plated sparsely on a 48 well plate and allowed to attach for 4 days, and then fixed in 4% paraformaldehyde and permeabilized with 0.1% triton. Cells were blocked in 5% BSA, stained with 1:200 cTnT primary antibody (Abcam:ab45932) and a 1:500 dilution of the 488 anti-rabbit antibody, stained with DAPI, and imaged on the Keyence microscope. Cell area was quantified with imageJ and Cell Profiler (21) and statistical differences in cell size were assessed with a non-parametric Mann-Whitney test in PRISM.

For assessment of α- and β-myosin expression, cells were fixed and permeabilized in 4% PFA and 0.05% triton for fifteen minutes, blocked in a goat serum solution for one hour, and incubated in 1:50 β myosin mouse primary antibody, 1:100 α myosin rabbit primary antibody overnight at 4°C. Secondary antibodies Goat anti mouse 488 and Goat anti rabbit 647 were used to visualize and quantify the proteins.

For electron microscopy, cells were grown on microcontact printed Aclar film for 1 week and the fixed with a mixture of 2% glutaraldehyde (EMS Ca# 16000) and 4% Formaldehyde (EMS Cat # 15700) in 0.1M Sodium Cacodylate (EMS Cat# 12300) pH 7.4 for 1 hr. The fixing solution was replaced with 1% Osmium tetroxide (EMS Cat# 19100) for 2 hours at room temperature, washed 3x with ultra-filtered water, then stained in 1% Uranyl Acetate at RT for 2 hours. Samples were dehydrated in a series of ethanol washes (50%, 70%, 95%, 100%) for 30 min. each. The samples were transferred to Propylene Oxide (PO) and then infiltrated with Embed-812 resin (EMS Cat#14120) with PO at 1:2, 1:1 and 2:1 ratios for 2 hours and then placed in resin for 2 hours and placed in 65°C oven overnight. Thin sections (80 nm) were cut along a plane parallel to the patterned substrate plane of the cell (near the plane of attachment) using a Leica EM UC7 Ultramicrotome and placed on 100 mesh coper grids (EMS). Samples were post-stained with 1.5% Uranyl Acetate in 50% Acetone for 30 seconds, then washed, dried, and then stained for 2 minutes in Sato's lead cirtrate and imaged with a JEM-1400 transmission electron microscope (JEOL) at 120 kV. Between 5 and 10 images were taken between 2500X and 3000X magnification with a on Gatan Orius 832 digital camera (Gatan, Pleasanton, CA) and sarcomere length and z-disk thickness was quantified in image J.

***Inhibitor study***

To clarify the effect of ERK and Akt, the cells were treated with either Akt inhibitor (Akti 1/2, Tocris: 5μM) or and ERK inhibitor (SCH772984, SelleckChem: 1μM) every 3 days between day 26 and day 46 (a time frame over which hiPSC-CMs have been shown to increase in area). At day 46, the cells were replated sparsely at a density of 50 thousand cells per coverslip and fixed and stained for β-myosin heavy chain and cardiac troponin T (Abcam:ab45932). The area of cells was quantified using Cell Profiler and the results from 2 batches of cells were combined and analyzed using Graphpad PRISM software with a 2-way ANOVA and the Kruskal-Wallis non-parametric test for multiple comparisons.

***Assessment of protein transcription, expression and activation***

Batch-matched cell lysates were collected from nine separate differentiations between day 50 and 60 of differentiation in either RIPA buffer supplemented with protease inhibitor, phosphatase inhibitor, and DNAse for western blot analysis or in TRIZOL LS for transcript analysis. For western blots, to measure both the activation of signaling proteins and the expression of myosin heavy chains, we used a 4-15% gradient gel (BIORAD) and separated at 150 V for 2 hours. Proteins were transferred to a nitrocellulose membrane at 80V for two hours on ice. These membranes were blocked in a 5% milk protein block solution in Tris buffered saline with 0.01% Tween (TBST), rinsed, and incubated overnight at 4 degrees in a 1:1000 solution of primary antibody in 2% BSA. After primary antibody incubation (pERK:Cell Signaling #9106S and pAkt: Cell Signaling #2965S), the gels were rinsed in TBST and incubated in 1:5000 secondary antibodies conjugated with HRP. After a final rinse, the gels were incubated for five minutes in Clarity ECL substrate (BIORAD), imaged on a chemidoc luminescent scanner, and band intensity was quantified in FIJI. After staining for the fluorescent protein, the membranes were stripped in a solution of 15 g glycine, 1 g SDS, 10 mL Tween 20 and distilled water (1L) and reblocked and reincubated in a 1:1000 dilution of primary for the total amount of protein (ERK:Cell Signaling 9102S and Akt: Cell Signaling #9272S). Many of these blots were also stained for α and β myosin antibodies that were developed by the Kraft lab and the Spudich lab. For each differentiation batch, the densitometry of measurements were normalized to the total protein, and then the sample from P710R cells were normalized to the control cells from that batch. Statistical significance was determined using a one sample t-test to test for a difference from 1.

***Computational modeling***

We used a previously validated model which incorporates discretized flux equations to describe transitions of myosin heads and thin filaments from inactive to active/bound states and predicts active and passive force (22, 23). This model also incorporates force-sensitive release of myosin heads from the thick filament and was modified to reflect the asymmetric load-dependent detachment rate (Figure S7). The fraction of binding sites in the active state was defined as $N_{on}$. The regulation of the thin filament activation and flux between myosin states were calculated according to the method described in a previous publication (23). $M_{OFF}$, $M_{ON}$, and $M_{FG,i}$ were defined as the fraction of myosin in the OFF state (equivalent to SRX), the ON state (available but unbound), and force generating states with i=1-41 different displacements (x) between the binding site and the zero load position of the myosin head. The range of potential positions of myosin heads relative to the no-load position ($x_i$) range from -10 to 10 nm with a bin size of 0.5 nm. A series of flux equations ($J_1$, $J_2$, $J_3$, and $J_4$) describe the transitions between these myosin states, and $J_1$, $J_2$, and $J_3$ were defined as previously described (23):

$$J_1 = k_{-SRX}(1 + k_{force} * F_{Total})M_{OFF}$$ Eqn. S1

$$J_2 = k_{+SRX} * M_{on}$$ Eqn. S2

$$J_3(x) = k_A * e^{\frac{-k_{cb}x^2}{2K_BT}} * M_{on}(N_{on} - N_{bound})$$ Eqn. S3

$J_1$ represents the flux of myosin heads from $M_{OFF}$ to $M_{ON}$ and is proportional to the fraction of heads in the SRX state. Based on previous modeling results and studies of the SRX state in muscle fibers and myofibrils, we assumed that the rate would be increased by increasing force in the muscle and so included a coefficient of force-sensitivity ($k_{force}$) with units of $N^{-1}$ $m^2$. $J_2$ represents the flux of myosin heads from $M_{ON}$ to $M_{OFF}$ and is proportional to the fraction of heads in the ON state. $J_3$ represents the flux of myosin heads from $M_{ON}$ to $M_{FG,i}$ and is proportional to $M_{ON}$ as well as the fraction of active, unbound binding sites on the thin filament. This flux is determined by the attachment rate of myosin to available actin ($k_A$). The Gaussian term reflects the probability of a myosin head with spring constant $k_{cb}$ being stretched to length x by thermal energy. $k_{cb}$ is the stiffness of the crossbridge spring and has units of N $m^{-1}$. $K_B$ is the Boltzmann's constant (1.38 x $10^{-23}$ J $K^{-1}$) and T is the temperature (set to 310 K). We modified the final flux equation $J_4$ describing bound myosins exiting the attached state to reflect the exponential load dependence observed in our single molecule measurements.

$$J_4(x) = k_0 * e^{\frac{-k_{cb}x\delta}{K_BT}} * M_{FG,i}(x_i)$$

if $|x_i| < 8$ nm, and

$$J_4(x) = (k_0 * e^{\frac{-k_{cb}x\delta}{K_BT}} + (|x_i| - 8) * p_{max}) * M_{FG,i}(x_i)$$ Eqn. S4

when $|x_i| > 8$ nm.

$k_0$ and $\delta$ were defined from HFS measurements and $p_{max}$ is the maximum parameter rate (set to 5000 $s^{-1}$). The overall rate of detachment increases towards $p_{max}$ as the displacement of the myosin head relative to its baseline position extends beyond 8 nm (Figure S8A) based on reported averages for powerstroke ranges and a natural limit for myosin head extension (24). Adding the muscle force (N $m^{-2}$) normalized by the number of myosin heads (1.2 x $10^{16}$ $m^{-2}$) into the calculation of detachment rate caused only a very small change in the overall fit (less than 10% increase in mean normalized deviation). The highest expected contribution of this normalized force at peak for these cells is less than half than the force of the individual cross bridge springs, so it was not included in these calculations. These flux expression were used to solve a series of differential equations at each time step (1 ms intervals) to calculate the fraction of myosin in each state and resultant total force, as previously described (23).

$$F_{total} = F_{active} + F_{passive}$$ Eqn. S5

$$F_{active} = N_0 k_{cb} \sum_{i=1}^{n} M_{FG,i}(x_i + x_{ps}) \quad \text{Eqn. S6}$$

$$F_{passive} = k_p(L - L_0) \quad \text{Eqn. S7}$$

The total force is comprised of active and passive force components. The active force depends on the powerstroke step size ($x_{ps}$), cross bridge stiffness ($k_{cb}$) and on the number of myosin heads in a hypothetical cardiac half-sarcomere with a cross-sectional area of 1 $m^2$ ($N_0$). $N_0$ was set to 1.2x $10^{16}$ for these simulations. This estimation was derived from the assumptions that there are ~100 myosin heads oriented towards actin per half thick filament (25), the half filaments have a spatial density of 4 x $10^{14}$ $m^{-2}$ (26), and the myofibrils in cell occupy ~30% of the cell space (by our observation in electron microscopy images). Multiplying these three numbers together (100 x 4 x $10^{14}$ x 0.3) gives an estimate of 1.2 x $10^{16}$ for the average myosin head density, but variation in the density of myofibrils or the density of heads on less mature thick filaments might contribute to the variability observed in the cells (Figure 7C). The passive force was determined based on a linear passive stiffness term ($k_p$), defined as 10 N $m^{-2}$ $nm^{-1}$, and the change in length from the baseline length ($L_0$) of 880 nm (to match the sarcomere lengths we measured in these cells). It also included a series stiffness of 100 N $m^{-2}$ $nm^{-1}$ that has previously been shown to improve model fit to relaxation dynamics (27). This parameter accounts for serial compliance of the cell and substrate and was adjusted to predict shortening of the sarcomeres (Figure S9E) within the range of cell shortening we measured for these cells (2-3%) and the percent sarcomere shortening measured in previous studies of hiPSC-CMs with MYH7 mutations (28).

To implement the modified model in the context of this project, we first incorporated the rates and parameters directly measured in our molecular characterization of the effects of the P710R mutation ($k_0$, $\delta$, and step size [$x_{ps}$]). We used previously reported actin activated ATPase measurements using WT and P710R sS1 constructs to define $k_A$ (29). The values were calculated from the previously published forward attachment rate of myosin ($k_A$) multiplied by the $K_{app}$ to account for changes in actin affinity and the fractional (unitless) description of actin availability in this model. This scaling of $k_A$ transforms it to units of $s^{-1}$ (compared with the reported units of $s^{-1}$ $\mu M^{-1}$ (29)). Finally, after calculating this rate constant ($k_A$) from previously reported values, we divided by the integral of the gaussian term (which depends on $k_{cb}$) to describe the density of flux at each segment of the available range of binding, making the final units calculated within the model ($s^{-1}$ $nm^{-1}$) consistent with rate constants previously reported using this model (23)). We also calculated the force at baseline ($F_{baseline}$) as the total force after 500 ms of equilibration (before initiation of the calcium transient) for each simulation.

We normalized our measured $Ca^{2+}$ traces and scaled them between pCa 6.17 and 6.72 according to the reported calibrated values from the previously published model simulations (23) as the input protocol for these simulations (Figure S9A). The rate constant for thin filament deactivation ($k_{off}$) was limited to 100-200 $s^{-1}$ and the $k_{+SRX}$ rate constant was held unchanged between control and mutant myosin simulations at 200 $s^{-1}$, consistent with the previous publication of this model (23). We initially set $k_{on}$, $k_{coop}$, $k_{force}$, $k_{cb}$ to previously reported values from cardiac tissue twitch and optimized these parameters to find the best fit of $F_{total} - F_{baseline}$ to our representative traces of cell forces normalized by cross sectional area. $k_{on}$ is a rate constant ($M^{-1}$ $s^{-1}$) that determines the rate of thin filament activation proportional to the concentration of $Ca^{2+}$. $k_{coop}$ is a unitless term that describes the interactions between regulatory units on the thin filament that contribute to myofilament cooperativity. These interactions further suppress thin filament activation at low $Ca^{2+}$ concentrations while augmenting activation at higher $Ca^{2+}$ concentrations. $k_{-SRX}$ for control was set to 20 $s^{-1}$, based on measurements of the SRX percentage in cell ranging from ~60 to 90% in hiPSC-CMs (28). After defining a parameter set that fit the WT curve, a specific optimization of the value for $k_{-SRX}$ for the P710R myosin was performed by initially setting the value of $k_{-SRX}$ to that of the WT myosin (20 $s^{-1}$) and allowed to varied between 0 and a 20 fold increase over WT. We also compared the fit to the data after optimizing the relative change in both $k_{-SRX}$ and $k_{force}$ or just $k_{+SRX}$ (Figure S9), but found less than 1% improvement in the mean normalized deviation (error term defined in previous description of the model (23)). We calculated confidence limits for these parameter fits by varying the parameter until the mean normalized deviation exceeded the best fit value by more than 5%, as previously described (23, 30). We also performed sensitivity analysis to determine the effect of each parameter on the active force and the %SRX at peak contraction (Figure S10). While holding the rest of the parameters constant at the best fit values, we varied one parameter to 0.1, 0.5, 2 and 10 times its original value and plotted the results.

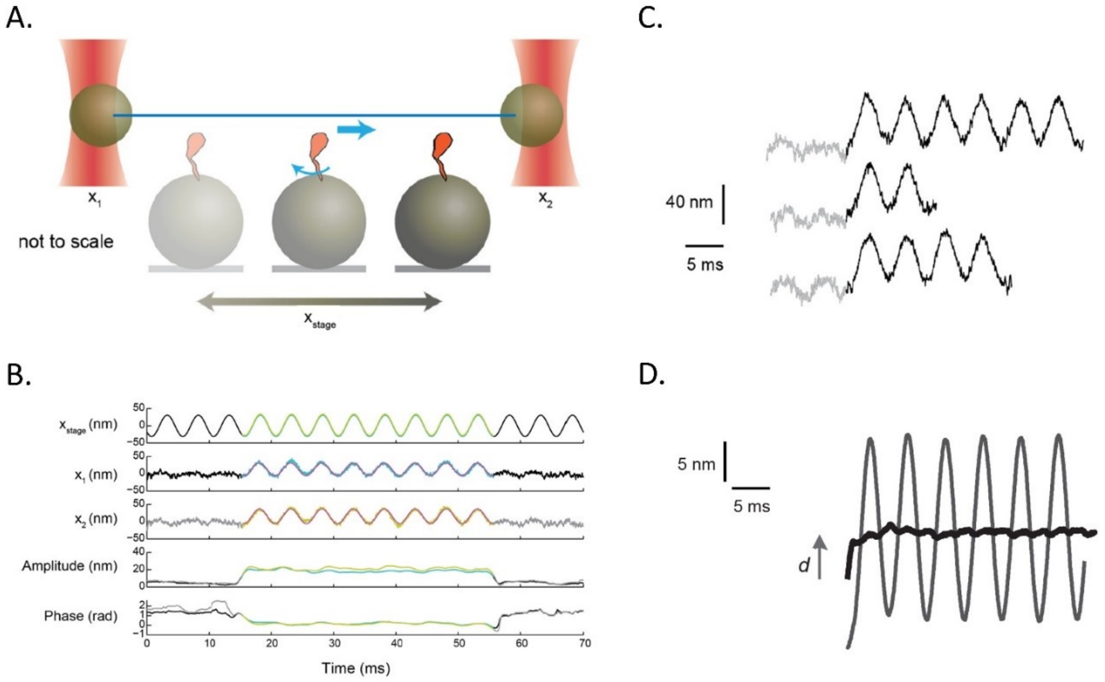


**Figure S1. Analysis of harmonic force spectroscopy (HFS) optical trap data to determine the step size of a single myosin molecule. A.** Schematic of HFS in the dual-beam optical trap. Myosin (orange) sits on top of a platform bead on a stage that oscillates sinusoidally with position $x_{stage}$. The positions of the trapped beads $x_1$ and $x_2$ are recorded. Upon binding of myosin to the actin (blue) stretched across the two trapped beads ("actin dumbbell"), myosin undergoes the power stroke, moving actin (blue arrow). **B.** The stage position $x_{stage}$, positions of trapped beads $x_1$ and $x_2$, amplitudes of $x_1$ and $x_2$, and phases of $x_1$ and $x_2$ relative to the stage position are shown around one example binding event. Upon myosin's attachment to actin, the amplitude of the sinusoidal oscillations in $x_1$ and $x_2$ increases while the phase decreases because the actin dumbbell is now strongly coupled to the oscillating stage. The detected binding event is highlighted in color. **C.** Time traces of three example events before (gray) and during (black) binding. **D.** All bound traces from one molecule (hundreds of binding events) are start-aligned, extended, and averaged (dark gray curve with large amplitude of oscillation). A fitted sine function is subtracted from the averaged trace, revealing the change in actin dumbbell position due to myosin binding alone (black). Myosin's power stroke is apparent within the first few milliseconds (gray arrow).

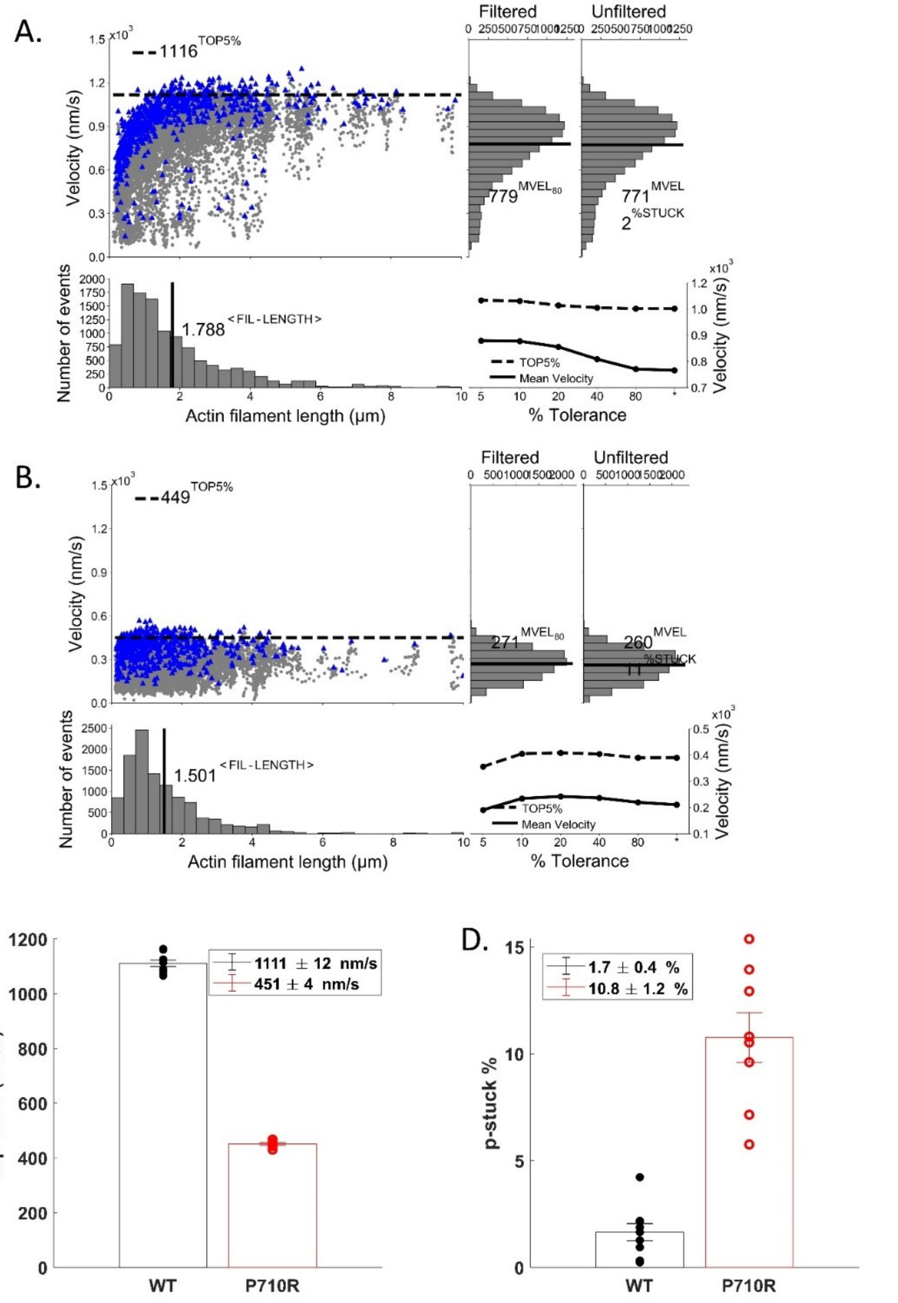


**Figure S2. Analysis of the actin-sliding velocities of P710R and WT sS1-AC in the unloaded motility assay. A.** and **B.** Fast Automated Spud Trekker (FAST) analysis of data from an example motility experiment using WT (A) (Movie S1) and P710R (B) (Movie S2) sS1-AC. The FAST parameters used for this paper were window size n = 5, path length p = 10, percent tolerance pt = 80, and minimum velocity for stuck classification minv = 20 nm/s. **C. and D.** Top 5% velocities (C) and percent stuck filaments (D) of all WT and P710R motility experiments. Each data point represents one experiment analyzed as in A and B. Values represent mean ± s.e.m. See also methods.

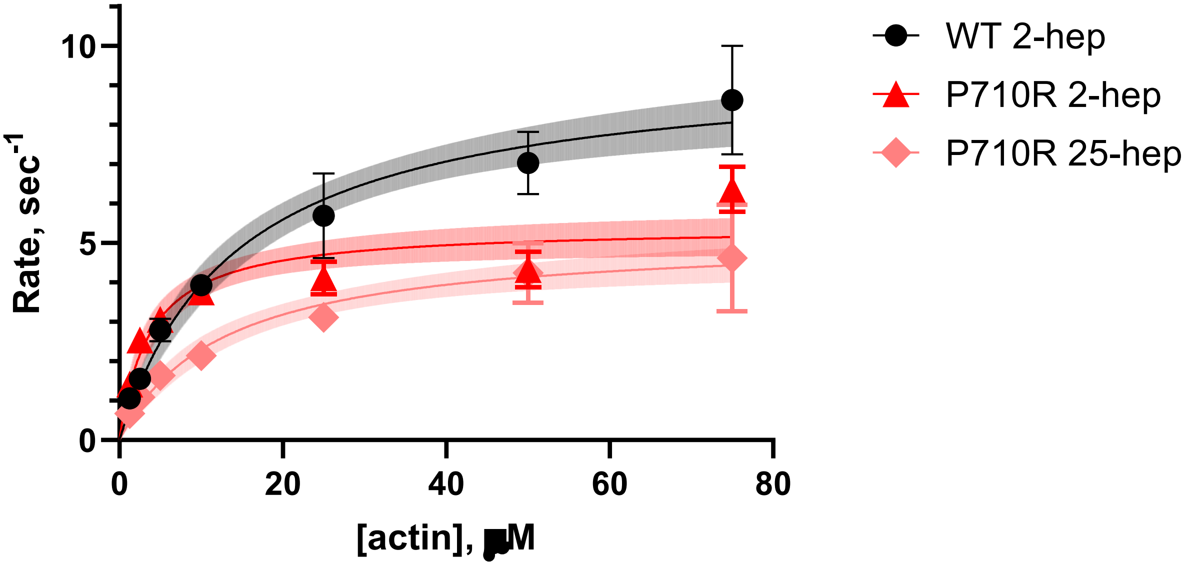


**Figure S3. ATPase for WT and P710R 2-hep and P710R 25-hep myosin.** Representative traces (3 replicates per concentration), one of two protein preps which gave similar results (and are presented as normalized results in Figure 3).

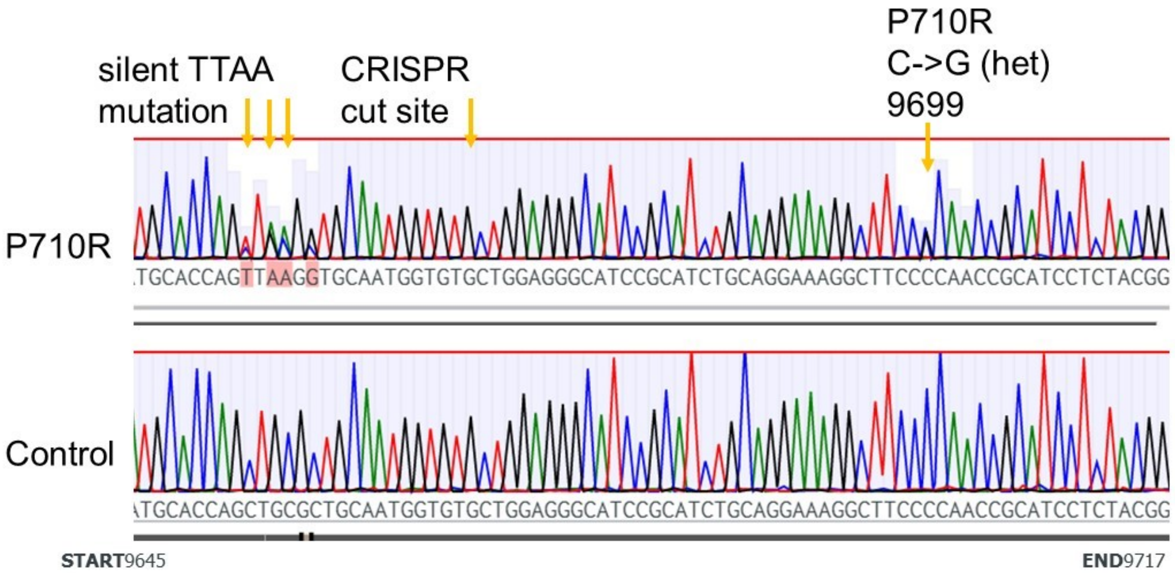


**Figure S4: Gene editing confirmation of heterogeneous editing at site of interest.** Excerpts of sequencing results of the MYH7 gene (between base pairs 9645 and 9717).

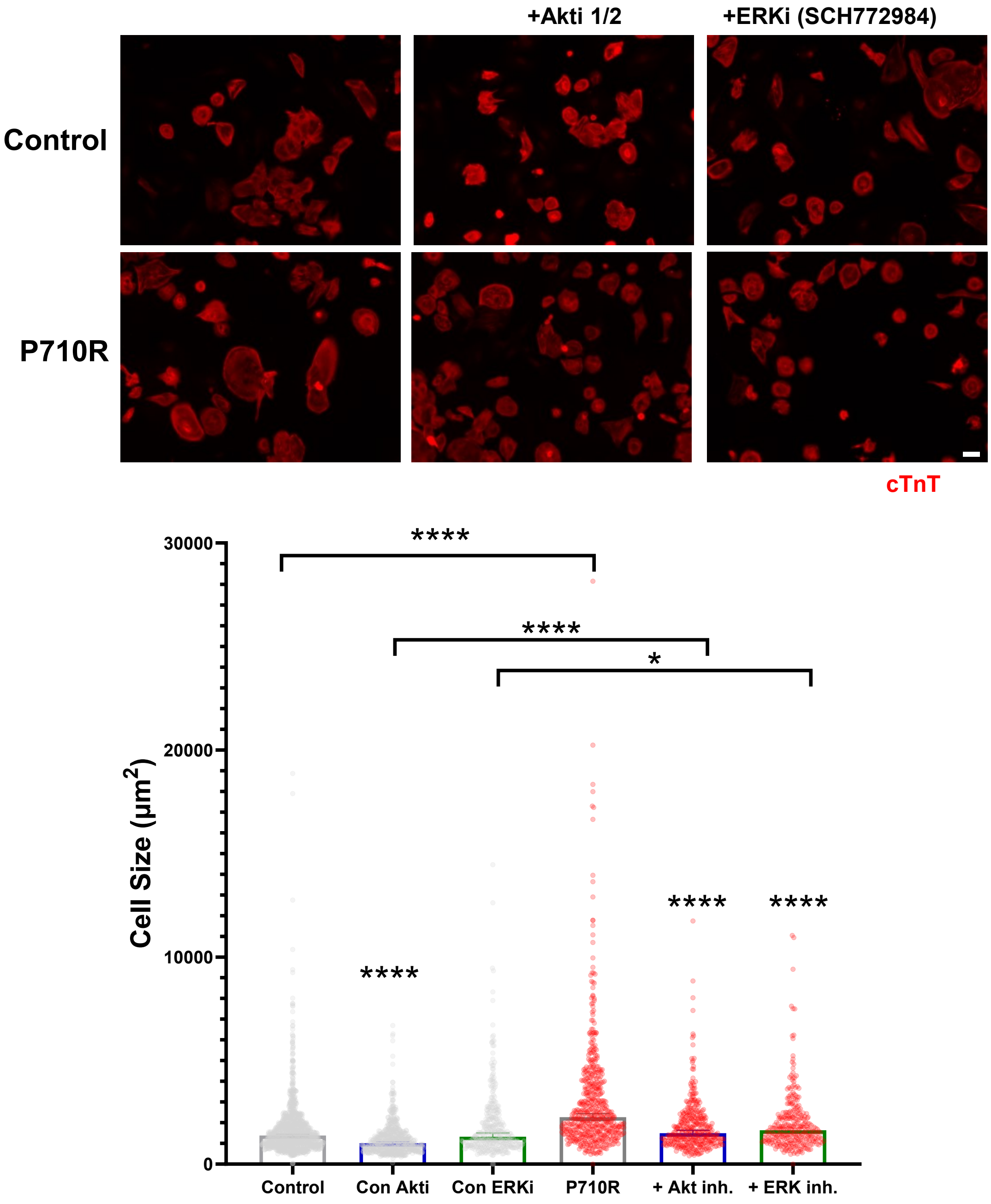


**Figure S5: Full analysis of effects of Akt and ERK inhibition.** Isogenic control cells were grown in parallel and treated with Akti and ERK inhibitors (Akti 1/2 and SCH772984) for 21 days (media change every 3 days). Scale bar represents 50 µm. The differences in size were quantified using Cell Profiler and analysis using a 2-way ANOVA and non-parametric test correcting for multiple comparisons. * signifies $p < 0.5$, **** signifies $p < 0.0001$.

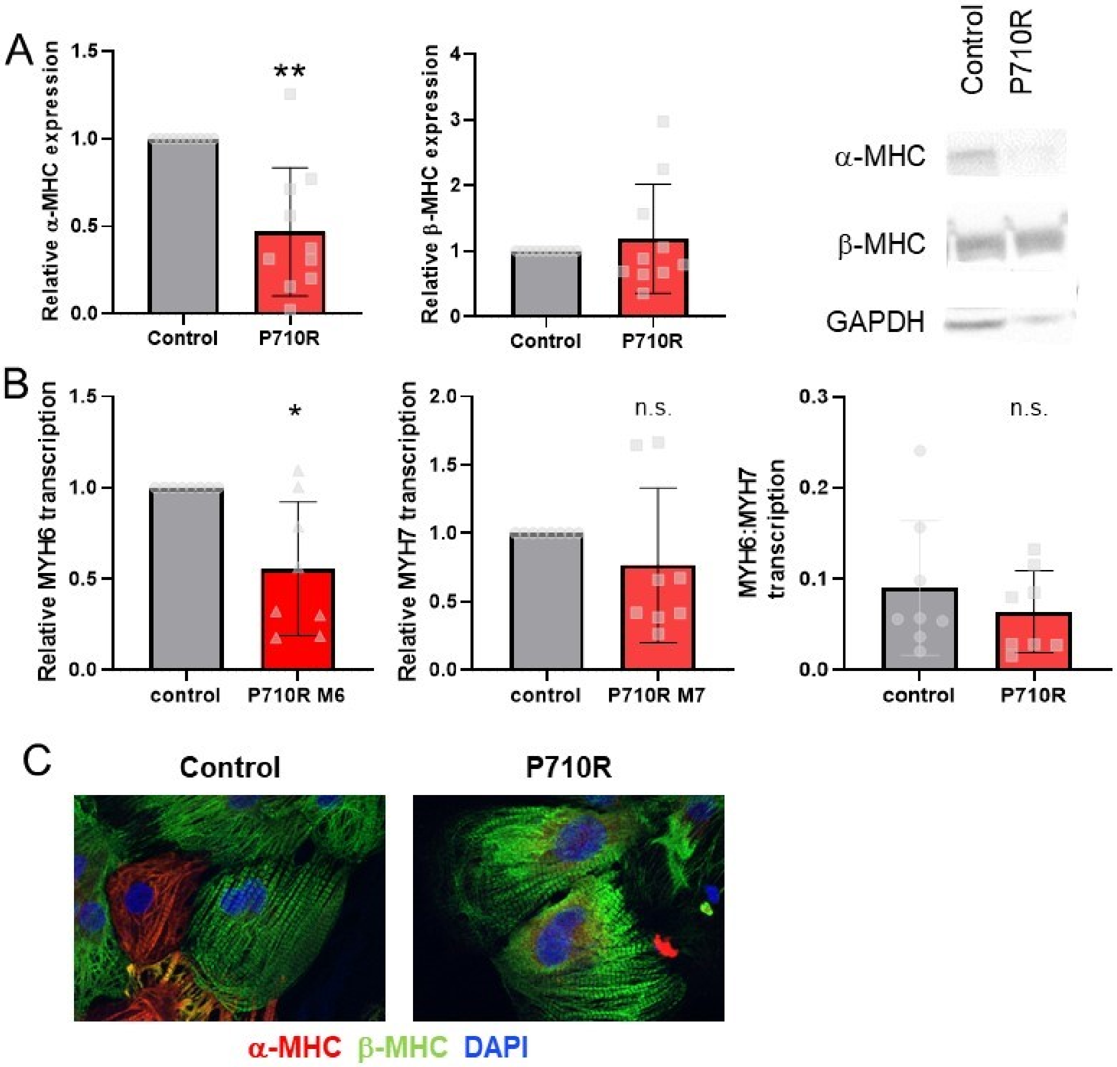


**Figure S6: Quantification of α and β myosin in cell population at day 45-50. (A)** Western blots were performed and the expression of each myosin type (quantified by densitometry and normalization against GAPDH) was further normalized to the protein expression level in the isogenic control cells for that differentiation batch. The mean relative expression was compared to 1 to test for significance (* indicates $p < 0.5$). (**B**) Relative transcription was also measured using qPCR. (**C**) Representative immunostaining images of unpatterned cells stained for α- and β-MHC.

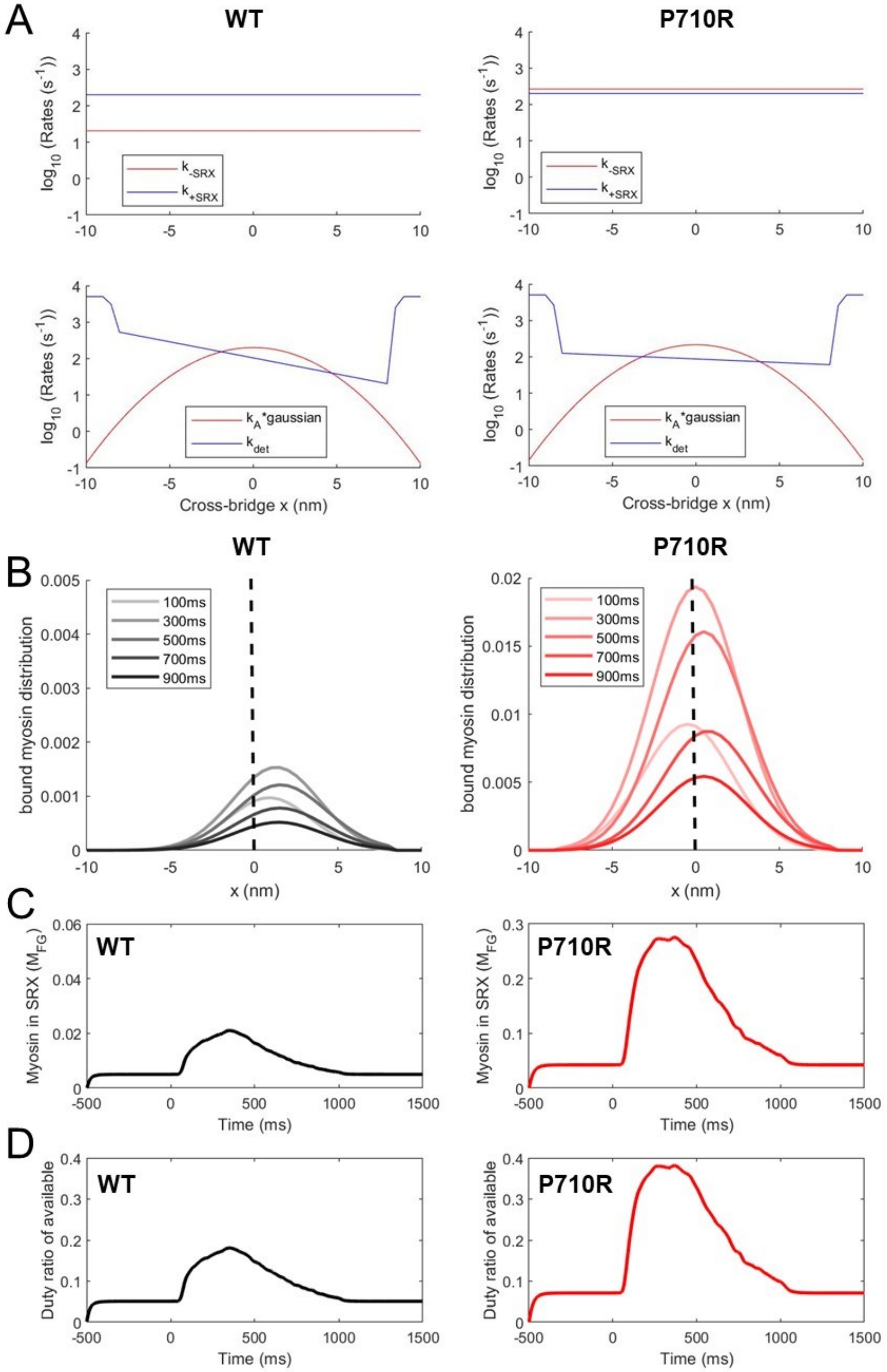


**Figure S7: Myosin transition rates for WT and P710R myosin influence crossbridge distribution.** (A) Transition rates after parameter estimation reflect the increase in $k_{-SRX}$ and decreased force sensitivity with the P710R mutation. (B) Analysis of the distributions of crossbridge lengths (x) during the contraction cycle show a more rightward shift (towards higher resistive loads) for the WT, with a more centered distribution for the P710R. (C) Attached, force generating (as a fraction of total) myosin over the time course of activation. (D) Calculated duty ratio as fraction of available myosin heads ($M_{ON} + \sum M_{FG}$).

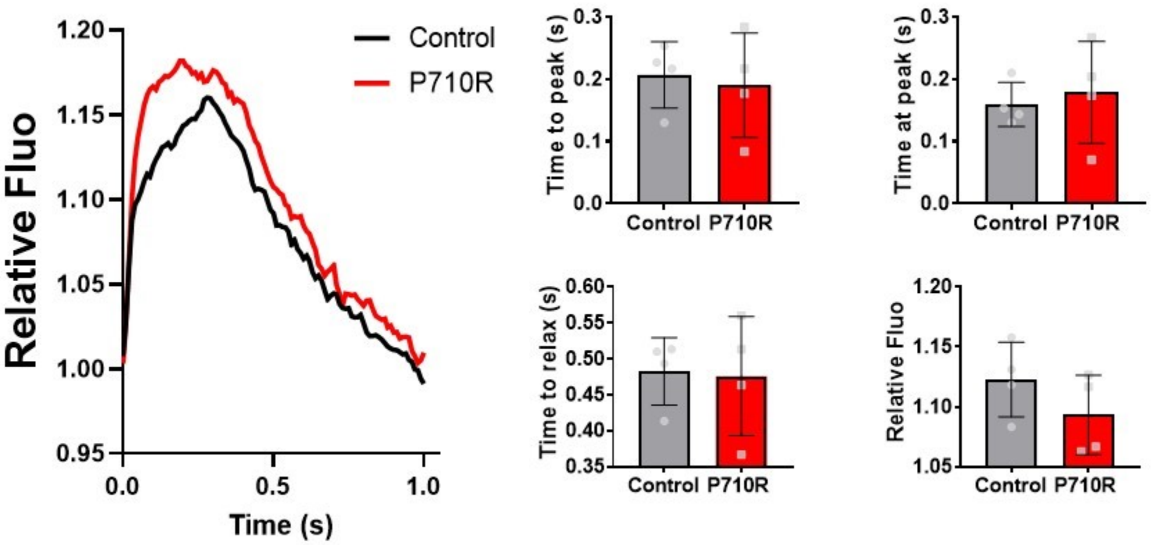


**Figure S8: Measurement of Calcium Transients in P710R cells.** (A) Representative transients from measurements of calcium in monolayers of hiPSC-CM. The time to peak was measured as the time between when the normalized signal crossed 5% of the range until it reached a threshold of 90% of the peak, and the time at peak was the time above a 90% threshold. Time to relax was quantified as the time to fall from 90% of peak to 5%, and the relative Fluorescence was the magnitude of peak fluorescence normalized to the baseline intensity. There were no significant differences between control and P710R cells.

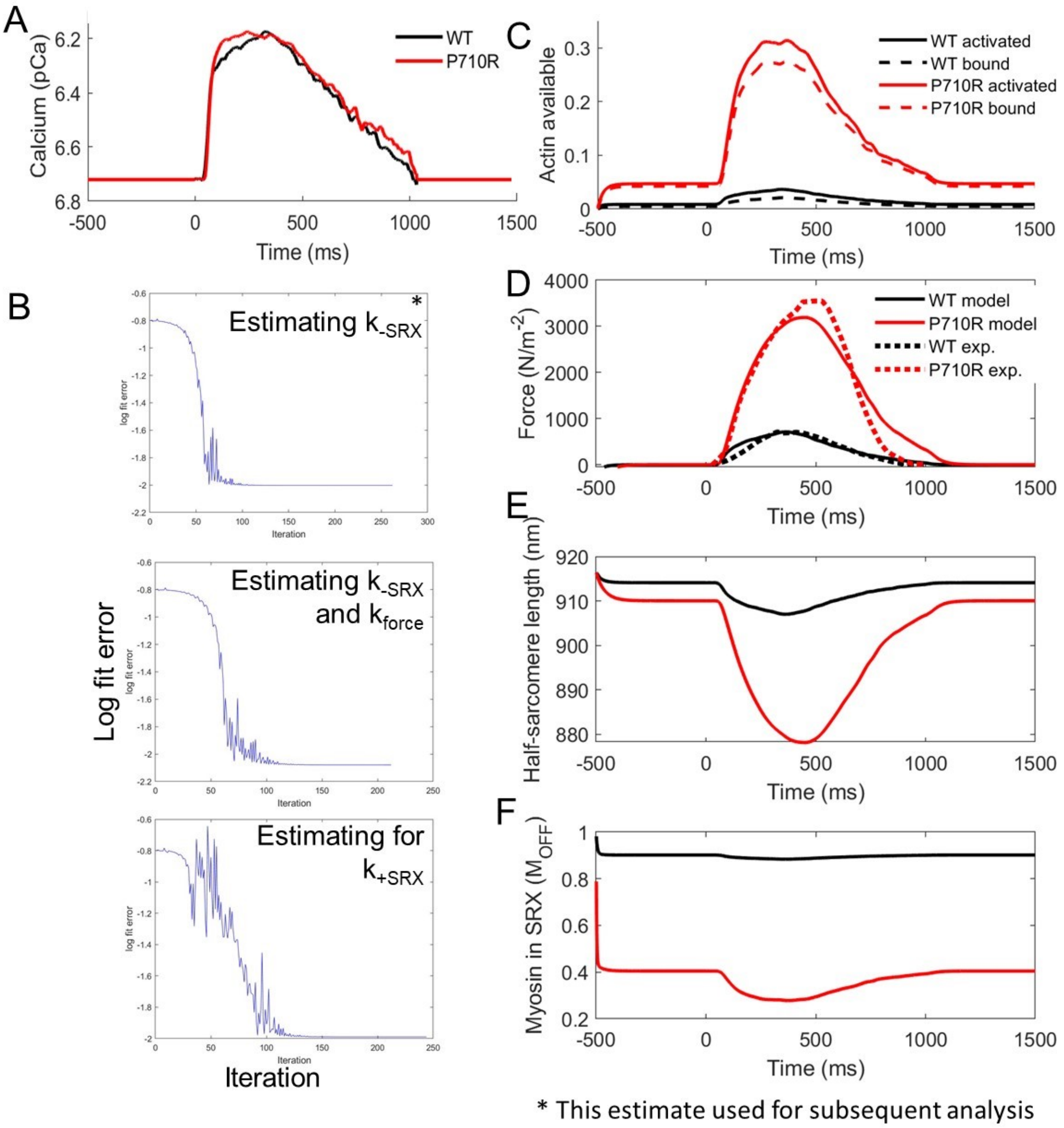


**Figure S9: Optimization and additional modeling predictions.** (A) Calcium transients measured in cellular monolayers were used as an input for the model. (B) After fitting parameters including $k_{force}$ and $k_{cb}$ to pedict the experimental control trace, optimization was used to find the best fit for $k_{-SRX}$, starting at the initial value of $k_{-SRX}$ for controls. (C) After fitting, the model predicted altered actin availability deriving from the changes in myosin kinetics. (D) The fitting technique was able to optimize the value of $k_{-SRX}$ for the best fit to the combined force data. Similar optimizations allowing both $k_{-SRX}$ and $k_{force}$ or only $k_{+SRX}$ to change followed a similar trajectory reducing error and ended with predictions similary to $k_{+SRX}$. (E) and (F) the predictions for sarcomere length change and SRX% qualitatively match what we and others have measured.

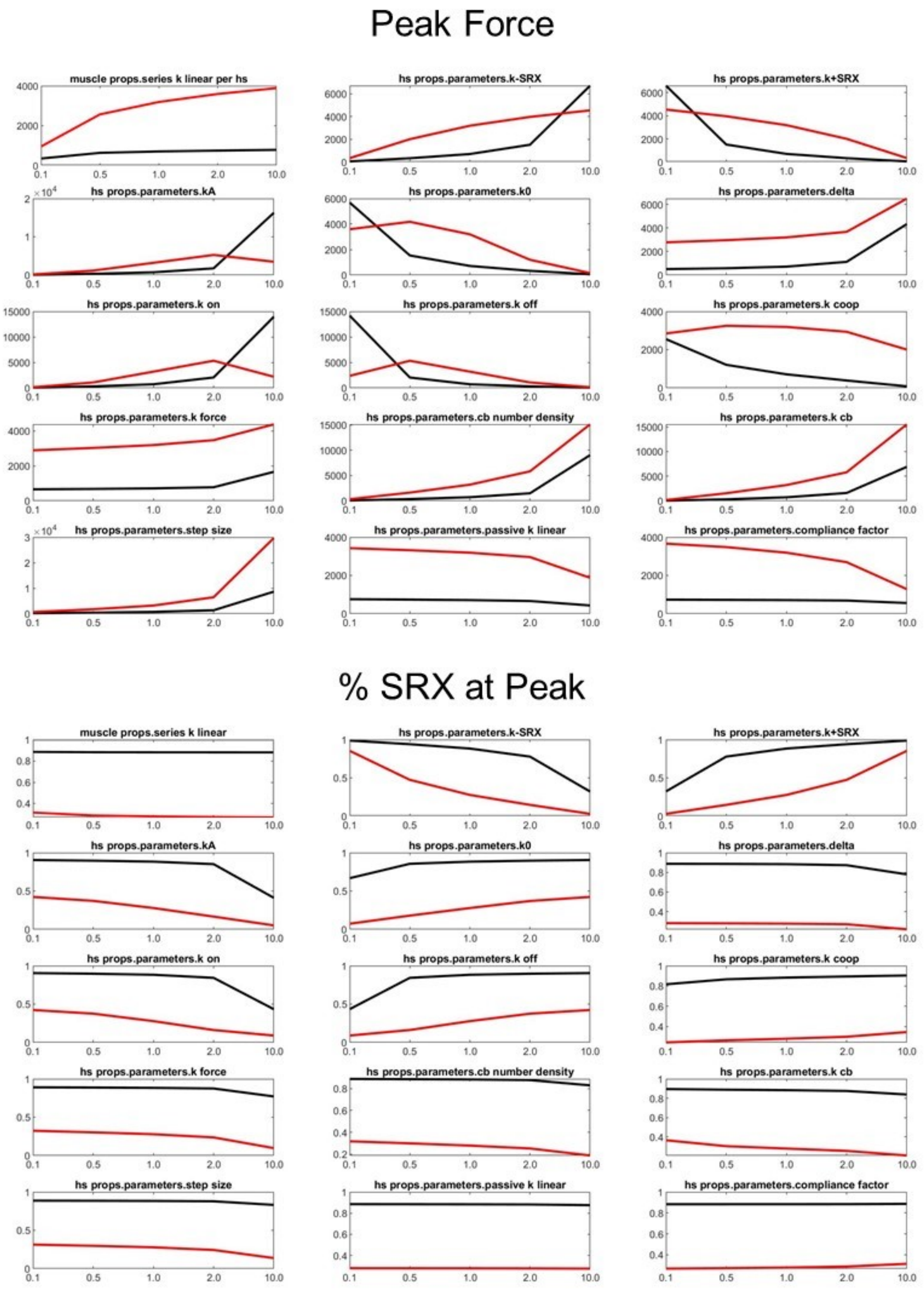


**Figure S10: Sensitivity analysis of individual parameter's effects on outputs.** Each parameter was varied from 0.1 to 10 times its initial value to show that parameters relative effects on peak force and SRX%.

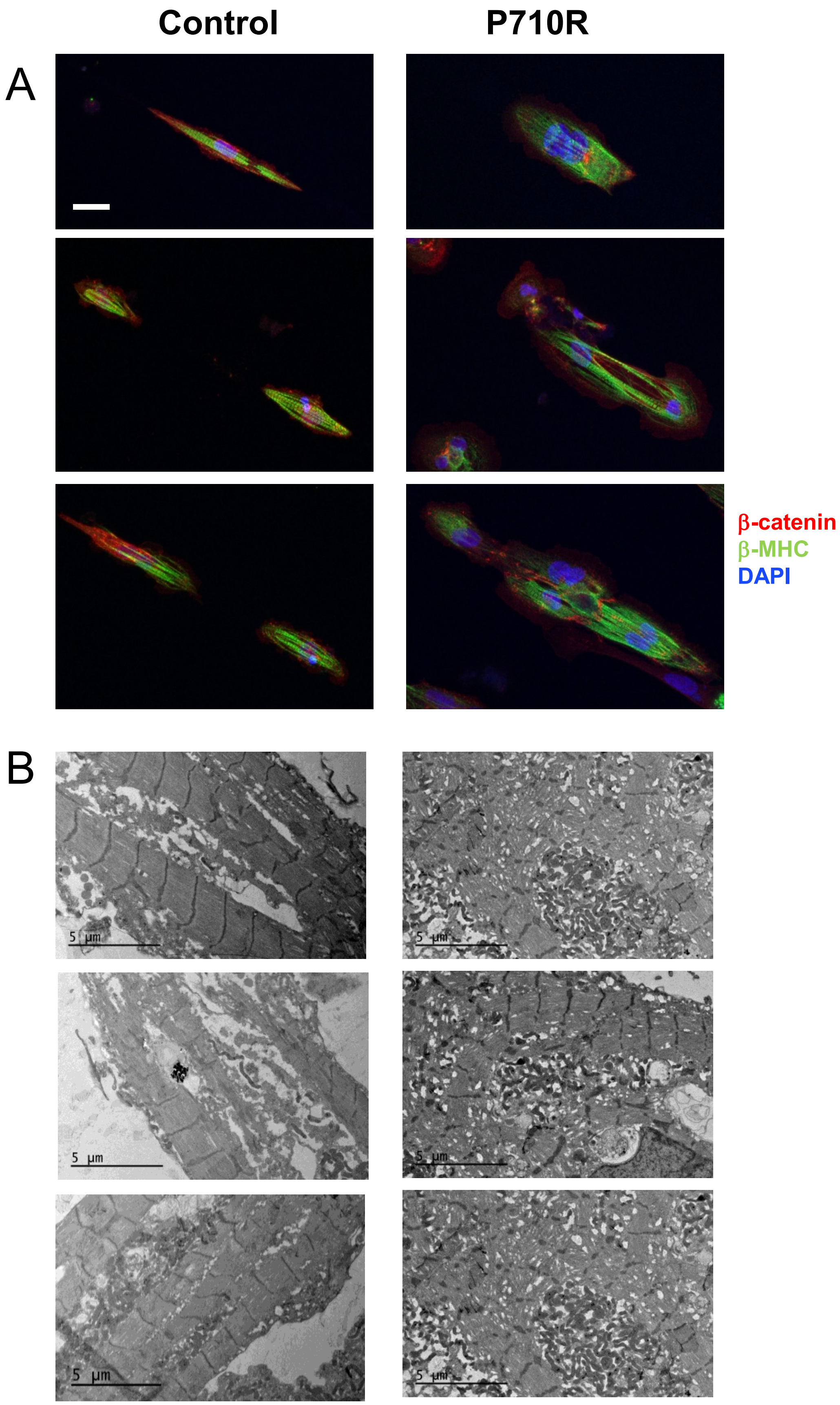
Control
P710R
A
B
β-catenin
β-MHC
DAPI
5 µm

**Figure S11: Additional myofibril distributions in micropatterned cells.** There is both variable density of myofibrils and variable structural maturity across cells when measured in (A) immunostained (Scale bar represents 20 μm) or (B) EM images.

**Table S1**. $k_{cat}$ and $K_M$ presented as fit (± SE of fit). $k_{fast}$, $k_{slow}$, and fast fraction are presented as mean ± SEM, and the text in blue is data previously published in Adhikari 2019 (15).

| | $k_{cat}$, $s^{-1}$ | $K_M$, μM | $k_{fast}$, $s^{-1}$ | $k_{slow}$, $s^{-1}$ | Fast fraction |
|---|---|---|---|---|---|
| WT 2-hep | 7.2 ± 0.7 | 12.8 ± 3.9 | 0.02 ± 0.002 | 0.002 ± 0.001 | 0.19 ± 0.03 |
| WT 25-hep | 4.1 ± 0.3 | 26.4 ± 5.1 | 0.02 ± 0.003 | 0.003 ± 0.001 | 0.59 ± 0.003 |
| P710R 2-hep | 4.5 ± 0.3 | 3.8 ± 1.1 | 0.04 ± 0.006 | 0.004 ± 0.001 | 0.18 ± 0.02 |
| P710R 25-hep | 4.3 ± 0.3 | 10.2 ± 2.5 | 0.024 ± 0.002 | 0.005 ± 0.001 | 0.27 ± 0.06 |

**Table S2: Parameters used in model after optimization.** Red parameter labels signify parameters which were different between control and mutant simulations, and asterisks signify the parameters which were optimized to find a best fit to the experimental force (with red asterisks denoting fitting of mutant parameter separately

| Parameter (units) | Control | P710R | Confidence limits for fits |
|---|---|---|---|
| $k_{-SRX}$ $(s^{-1})$ | 20 | 258* | [215-292] |
| $k_{force}$ $(N^{-1}\ m^2)$ | 6E-5 | 6E-5 | |
| $k_{+SRX}$ $(s^{-1})$ | 200 | 200 | |
| $k_A (s^{-1})$ | 656.7 | 701.5 | |
| $k_0$ $(s^{-1})$ | 104 | 87 | |
| $\delta$ $(nm)$ | 1.39 | 0.3 | |
| $x_{ps}$ $(nm)$ | 5.2 | 1.9 | |
| $k_{cb}$ $(N\ m^{-1})$ | 0.00063* | 0.00063* | [0.00058-0.00064] |
| $k_{on}$ $(M^{-1}\ s^{-1})$ | 3.2E7* | 3.2E7* | [3.02E7 - 4.0E7] |
| $k_{off}$ $(s^{-1})$ | 200 | 200 | |
| $k_{coop}$ *(unitless)* | 7* | 7* | [6.8-8.1] |
| $k_p$ $(N\ m^{-2}\ nm^{-1})$ | 10 | 10 | |
| $N_0$ $(\#\ m^{-2})$ | 1.2E16 | 1.2E16 | |

**Supplemental Table 3:** Summary results of output forces and SRX percentages with single variable perturbations

| *Condition* | *Actin availability* | *Peak Force* | *Baseline Force* | *%SRX at peak* | Baseline %SRX |
|---|---|---|---|---|---|
| *Optimized fit control* | 0.037 | 711 | 589 | 88.4 | 90.2 |
| *Optimized fit P710R* | 0.316 | 3193 | 998 | 27.8 | 40.4 |
| *Only $k_{-SRX}$* | 0.25 | 770 | 2101 | 25.8 | 39.2 |
| *Only kA* | 0.039 | 338 | 606 | 88.2 | 90.1 |
| *Only $x_{ps}$* | 0.036 | 644 | 473 | 88.7 | 90.2 |
| *Only $k_{det}$* | 0.038 | 263 | 563 | 88.3 | 90.2 |
| *Not $k_{-SRX}$* | 0.039 | 2821 | 447 | 88.4 | 90.2 |
| *Not kA* | 0.290 | 8660 | 947 | 29.1 | 40.7 |
| *Not $x_{ps}$* | 0.347 | 3872 | 2041 | 22.6 | 39.0 |
| *Not $k_{det}$* | 0.275 | 3062 | 1260 | 28.0 | 40.2 |

**Movie S1 (separate file).** Actin motility of WT β-cardiac myosin sS1-AC. Playback speed 3.5x.

**Movie S2 (separate file).** Actin motility of P710R β-cardiac myosin sS1-AC. Playback speed 3.5x.
